# Mycobacterium abscessus virulence traits unraveled by transcriptomic profiling in amoeba and macrophages

**DOI:** 10.1101/529057

**Authors:** Violaine Dubois, Alexandre Pawlik, Anouchka Bories, Vincent Le Moigne, Odile Sismeiro, Rachel Legendre, Hugo Varet, Pilar Rodriguez, Jean-Louis Gaillard, Jean-Yves Coppée, Roland Brosch, Jean-Louis Herrmann, Fabienne Girard-Misguich

## Abstract

Free-living amoebae might represent an evolutionary niche. In order to get more insights into the potential amoebal training ground for *Mycobacterium abscessus*, we characterized its full transcriptome in amoeba (Ac) and macrophages (Mφ), as well as the *Mycobacterium chelonae* intra-Ac transcriptome for comparison. Up-regulated genes in Ac allowed *M. abscessus* to resist environmental stress and induce defense mechanisms, as well as showing switch from carbohydrate carbon sources to fatty acid metabolism. Eleven genes implicated in the adaptation to intracellular stress, were mutated, with all but one confirmed to be involved in *M. abscessus* intra-Mφ survival. Cloning two of these genes in *M. chelonae* increased its intra-Mφ survival. One mutant was particularly attenuated in Mφ that corresponded to the deletion of an Eis N-acetyl transferase protein (*MAB_4532c*). Taken together, *M. abscessus* transcriptomes revealed the intracellular lifestyle of the mycobacteria, with Ac largely contributing to the enhancement of *M. abscessus* intra-Mφ survival.

## Introduction

To date, most of the known mycobacterial species are environmental organisms found in soil (Lavania et al., 2008), air (Angenent et al., 2005) and water (Ben Salah et al., 2009; Gomila et al., 2008; Guidotti et al., 2008), and belong to the Rapid Growing Mycobacteria (RGM). In contrast, most of the host-associated mycobacteria are slow growing mycobacteria (SGM). However, some exceptions exist. *Mycobacterium abscessus* and members of the *Mycobacterium chelonae* complex, in addition to *Mycobacterium fortuitum*, represent the main opportunistic pathogens among RGM (Griffith et al., 2007).

Identification of the last common ancestor of the genus *Mycobacterium* is still a matter of debate, although recent studies proposed the ancestor as an environmental species that evolved either as a soil-borne mycobacterium, as a waterborne mycobacterium such as *Mycobacterium avium-intracellulare* and *Mycobacterium marinum*, or as a host-associated mycobacterium such as *Mycobacterium avium* subsp *paratuberculosis* and *M. avium* subsp *avium*, *Mycobacterium tuberculosis* complex (MTBC) species and *Mycobacterium leprae* (Ahmed et al., 2007; Stinear et al., 2008).

Compared to other non-tuberculous mycobacteria (NTM), recovery of *M. abscessus* from the environment is rare (Thomson et al., 2013). Information from its genome sequence indicates its presence at the interface of soil, vegetation and water, an environment where free-living amoebae (FLA) are commonly found (Ripoll et al., 2009). FLA have been isolated from habitats in common with mycobacteria (Falkinham, 2009; Thomas and McDonnell, 2007) including cold-drinking water systems (Eddyani et al., 2008; Thomas et al., 2006), hot water systems in hospitals and cooling towers (Pagnier et al., 2008). FLA are ubiquitous organisms that feed on bacteria, and these bacteria have likely developed adaptations to the intracellular lifestyle to become amoeba-resistant bacteria (ARB) (Adékambi *et al.*, 2006; Lamrabet *et al.*, 2012). Mycobacteria have been isolated from such habitats by amoebal enrichment (Thomas et al., 2006)(White et al., 2010), allowing potential evolution toward pathogenicity by the acquisition of virulence genes by horizontal gene transfer (Gutierrez *et al.*, 2005; Ripoll *et al.*, 2009). Finally, amoebae are often considered as an ancestral form of macrophages (Mφ) sharing similar cellular structures and biological features (Barker and Brown, 1994; Greub and Raoult, 2004; Siddiqui and Khan, 2012).

*M. abscessus* has been shown to be resistant to amoeba phagocytosis and encystment, a property shared with all mycobacteria with the exception of the attenuated *M. bovis* BCG vaccine strain (Adékambi *et al.*, 2006; Salah *et al.*, 2009; Bakala N’Goma *et al.*, 2015). In addition, co-culture of *M. abscessus* with *Acanthamoeba castellanii* (Ac) increases its virulence when aerosolized in mice (Bakala N’Goma *et al.*, 2015) suggesting the existence of an amoebal ‘training ground’ for opportunistic pathogenic mycobacteria. Similarly, co-culture of amoebae with *M. avium* was found to trigger *M. avium* virulence by enhancing both entry and intracellular multiplication of the bacterium (Cirillo et al., 1997). The essential role of the ESX-4 *M. abscessus* type VII secretion system (T7SS) has also been demonstrated based on an intra-amoebal viability screen of *M. abscessus*, unraveling for the first time the active role of ESX-4 in intracellular *M. abscessus* survival (Laencina et al., 2018).

In order to gain more insights into the proposed amoebal training ground for *M. abscessus*, we characterized the full transcriptome of *M. abscessus* in Ac and Mφ, as well as the *M. chelonae* intra-amoebal transcriptome for comparison and characterization of *M. abscessus* virulence adaptations to intracellular life. Although amoebae and Mφ share common features, it has been shown that an intra-amoebal life requires specific adaptations (Laencina et al., 2018). A full description and analysis of *M. abscessus* transcriptomes will allow the identification of essential genes and a complete picture of *M. abscessus* intracellular replication and survival, whether in an amoeba environmental host or in Mφ, and elucidation of the mechanisms employed by the bacteria to resist host responses.

## Results

### Overall description of the *M. abscessus* intracellular transcriptomes

RNAseq data from *M. abscessus* planktonic or intracellular cultures (3 to 4 replicates per condition) were analyzed and compared to identify *M. abscessus* genes that were up- or down-regulated after Ac and Mφ co-cultures. Transcriptomes of *M. abscessus* in-Ac 4 and 16 hours post-infection (hpi), in Mφ 16 hpi and transcriptomes of *M. chelonae* in Ac 16 hpi were obtained and invariant genes were excluded from the analyses. Normalization and hierarchical clustering of normalized raw date confirmed the quality of transcriptomes (Sup. **Figure 1**). Differentially expressed genes (DEGs) were identified using the *DESeq2* package (Love et al., 2014) (Sup. Table 1) and the Log_2_ fold change (FC) values from *M. abscessus* transcriptomes were compared (**Sup. Figure 2**). In Ac, most DEGs up-regulated or down-regulated at 4hpi were still up- or down-regulated at 16hpi (**Figure 1A**). In order to detail the biological changes that correlated with *M. abscessus* intracellular regulation, we first grouped the DEGs in cluster of orthologues (Cluster of Orthologous Groups or COGs) (Tatusov et al., 2000). Highly up-regulated genes in Ac were more frequently found in COG O (Post-translational modification, protein turnover, chaperones), COG K (Transcription) and COG I (Lipid transport and metabolism) compared to the genome reference, although this last category tended to be under-represented at 16 hpi (**Figure 1B**). Comparatively, highly down-regulated genes were assigned to COG E and COG F (amino-acid and nucleotide transport and metabolism respectively) (**Figure 1B**).

**Figure 1:**
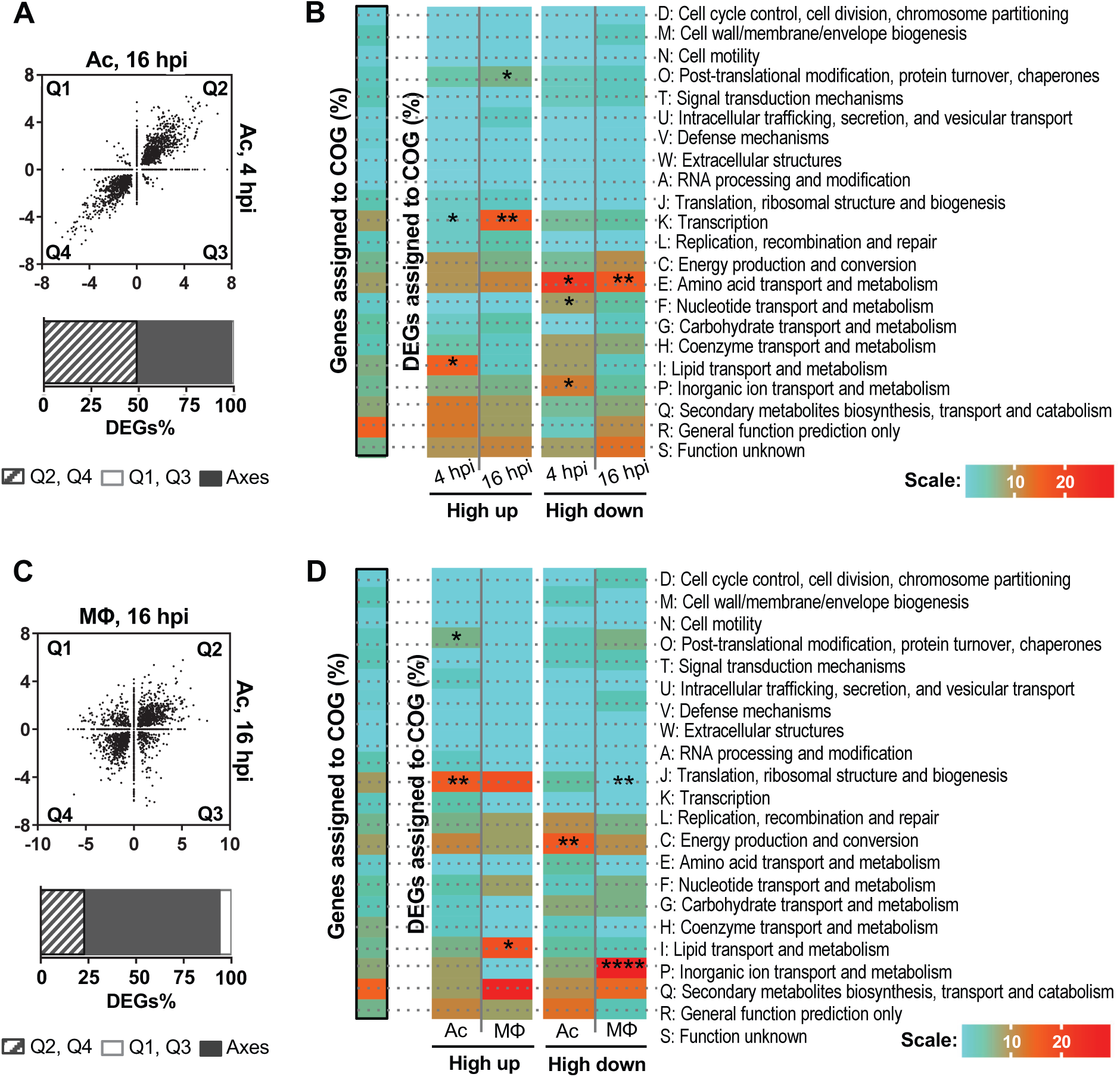
Description of *M. abscessus* transcriptomes in Ac (A) and Mφ (C). Differentially expressed genes (DEGs) fold change (FC) in *A. castellanii* (Ac) 4 hpi are plotted against DEGs FC 16 hpi (A). DEGs FC in Ac 16 hpi are also plotted against DEGs FC in Macrophages (Mφ) 16 hpi (C). DEGs froM quadrant 2 (Q2) and 4 (Q4) are genes regulated in the same direction whereas DEGs froM Q1 and Q3 are genes regulated in opposite direction. Dots on the plot axes are genes regulated in one condition only. Proportions of DEGs in each case are quantified. *M. abscessus* adaptations to Ac (B) and Mφ (D) unraveled by COG categorization Highly regulated genes were assigned to COGs. The genome assignation (framed in black) serves as a reference for COG enrichment tests. Fisher’s exact tests were performed to compare the transcriptome sets and the genome set of gene assignments to COG. * *p*<0.05. ** *p*<0.01. *** *p*<0.001. **** *p*0.0001.

When comparing *M. abscessus* Ac-16hpi *vs*. Mφ-16hpi transcriptomes, DEGs were found more dispersed, with 20% of DEGs regulated in the opposite direction (increased versus decreased and vice versa) (**Figure 1C**). The representation of DEG according to their FC highlighted that in Ac and Mφ most DEGs showed low changes (FC<|2|) 16 hpi (**Sup. Figure 2A**). In Ac, UP regulated DEGs predominated in comparison with their behavior in a Mφ environment (**Sup. Figure 2B**).

COG designations highlighted the differences between *M. abscessus* high DEGs in Mφ and Ac (**Figure 1D**). COG O, which was over-represented in the *M. abscessus* highly up-regulated genes in Ac, was more frequently associated with highly down-regulated genes in Mφ (**Figure 1B-D**). By comparison, COG P (Inorganic ion transport and metabolism) was over-represented in the *M*. *abscessus* transcriptome in Mφ only, potentially illustrating different adaptations to amoebal and Mφ environments (**Figure 1D**).

### Main biological pathway changes of *M. abscessus* in Ac and Mφ

We performed a gene ontology enrichment (GOE) analysis to further characterize the *M. abscessus* adaptions in Ac and Mφ. GOE were qualified by an enrichment factor (EF) (1 to 4) and a number of significantly enriched genes (from small (<10) to large (>100)) (**Figure 2**). In Ac, the most enriched up-regulated *M. abscessus* genes fell into polyamine transport (GO:0015846) to adenine salvage (GO:0006168), including small groups of genes involved in sulfur metabolism (sulfate assimilation pathway (SAP) (GO:0000103), hydrogen sulfide (H2S) biosynthetic pathway (GO:0070814) and detoxification (iron-sulfur cluster assembly (GO:0016226) (**Figure 2A**).

**Figure 2:**
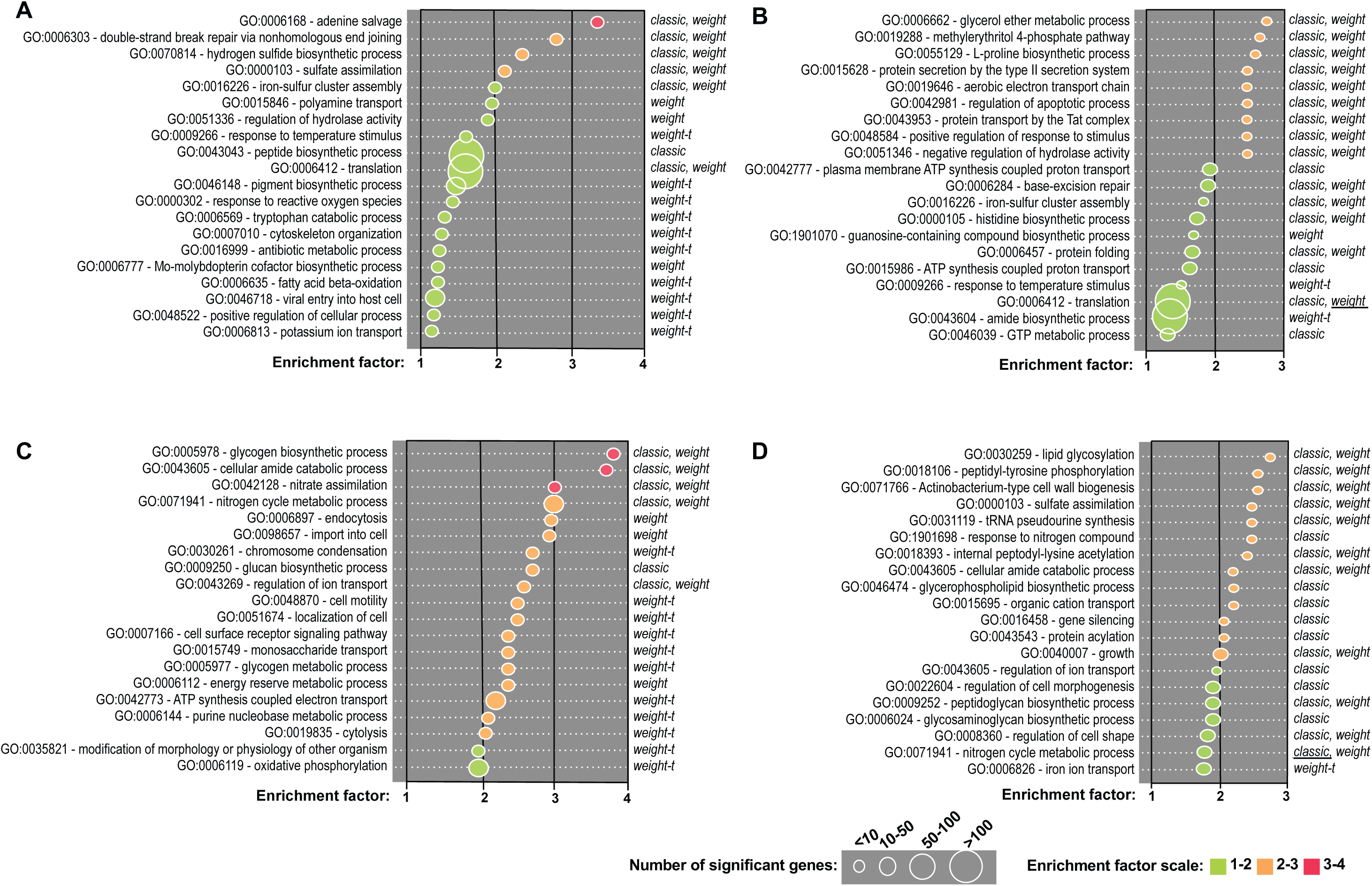
Gene ontology enrichment analyses applied on *M. abscessus* intracellular transcriptomes. **A**. Up-regulated genes in Ac 16 hpi. **B**. Up-regulated genes in Mφ. **C**. Down-regulated genes in Ac 16 hpi. **D**. Down-regulated genes in Mφ 16 hpi. Gene onthology (GO) enrichment analysis was performed with the topGO R package. Enriched GOs are sorted according to their enrichment factor (EF), corresponding to the ratio of significant DEGs assigned to the GO over expected assigned DEGs to the GO. Enriched GOs are represented by circles which size is proportional to the amount of significant DEGs assigned. Positive statistical tests are given that face each GO. Method giving the smallest *p*-value is underlined.

In Mφ, *M. abscessus* up-regulated enriched genes fall into different GO in comparison to those up-regulated in Ac. L-proline biosynthetic process (GO:0055129); methylerythritol 4-phosphate MEP pathway (GO:0019288) and Glycerol ether metabolic process (GO:0006662) were the most enriched (**Figure 2B**). These GO are followed by the type II secretion system and notably the Tat (twin-arginine translocation) pathway.

In Ac *M. abscessus* infections, the most enriched down-regulated genes fell into the nitrate assimilation GO (GO:0042128), to glycogen biosynthetic process GO (GO:0005978) (**Figure 2C**). In particular GO related to transport and metabolism of glucose were enriched (GO:0005977, GO:00015749, GO:0009250, GO:0005978) (**Figure 2C**).

In Mφ, *M. abscessus* down-regulated genes related to growth and parietal activities. From GO:0040007 corresponding to growth, up to GO:0030259 corresponding to lipid glycosylation, in addition to GO:0071941 (nitrogen cycle metabolism process), GO:0009259 (peptidoglycan synthesis) and GO:0022604 (regulation of cell morphogenesis), GOE indicated that *M*. *abscessus* slows down its energy-lycost metabolic processes and growth rate (**Figure 2D**). Taken together, these observations suggest that *M. abscessus* enters a slow-replicative state in Mφ and dedicates its energy to detoxification and protein secretion into the host.

### Regulation of the central carbon Metabolism of *M. abscessus* in Ac and Mφ

Following the GOE analysis, we investigated the different *M. abscessus* up- and down-regulated genes from metabolic pathways in Ac or Mφ. The major finding was that *M. abscessus* switches from a simple sugar-based carbon source to fatty acids inside Ac and Mφ (**Figure 3**). The glycolysis/neoglucogenesis and pentose phosphate pathways were mostly down-regulated or unchanged inside cells whereas the β-oxidation of fatty acids was up-regulated in Ac and Mφ. This switch was observed from the early time points after Ac infection. Fifteen genes predicted to encode enzymes necessary for the biochemical activation and β-oxidation of fatty acids were up-regulated in Ac and Mφ, such as: fatty acid-coenzyme A (CoA) synthase (*fadD3*, *9*, *10*, *19*); acyl-CoA dehydrogenase (*fadE5*, *14*, *23*-*24*, *27*-*29*, *31*); enoyl-CoA hydratase (*echA19*); hydroxy-butyryl-CoA dehydrogenase (*fadB2*) and acetyl-CoA transferase (*fadA5*, *6*). Genes implicated in the synthesis of enzymes involved in the breakdown of cholesterol A and B rings were highly induced in Ac and Mφ.β-oxidation of fatty acid and cholesterol breakdown result in the accumulation of propionyl-CoA that is detoxified by the methylmalonyl pathway. By-products of these 3 pathways and the GABA shunt feed the TCA cycle. The succinate generated by the TCA cycle enables the bacterium to deal with anaerobic respiration (Hartman et al., 2014). In addition, *M. abscessus* may detoxify glyoxylate by converting it into malate via the glyoxylate shunt (**Figure 3**).

**Figure 3:**
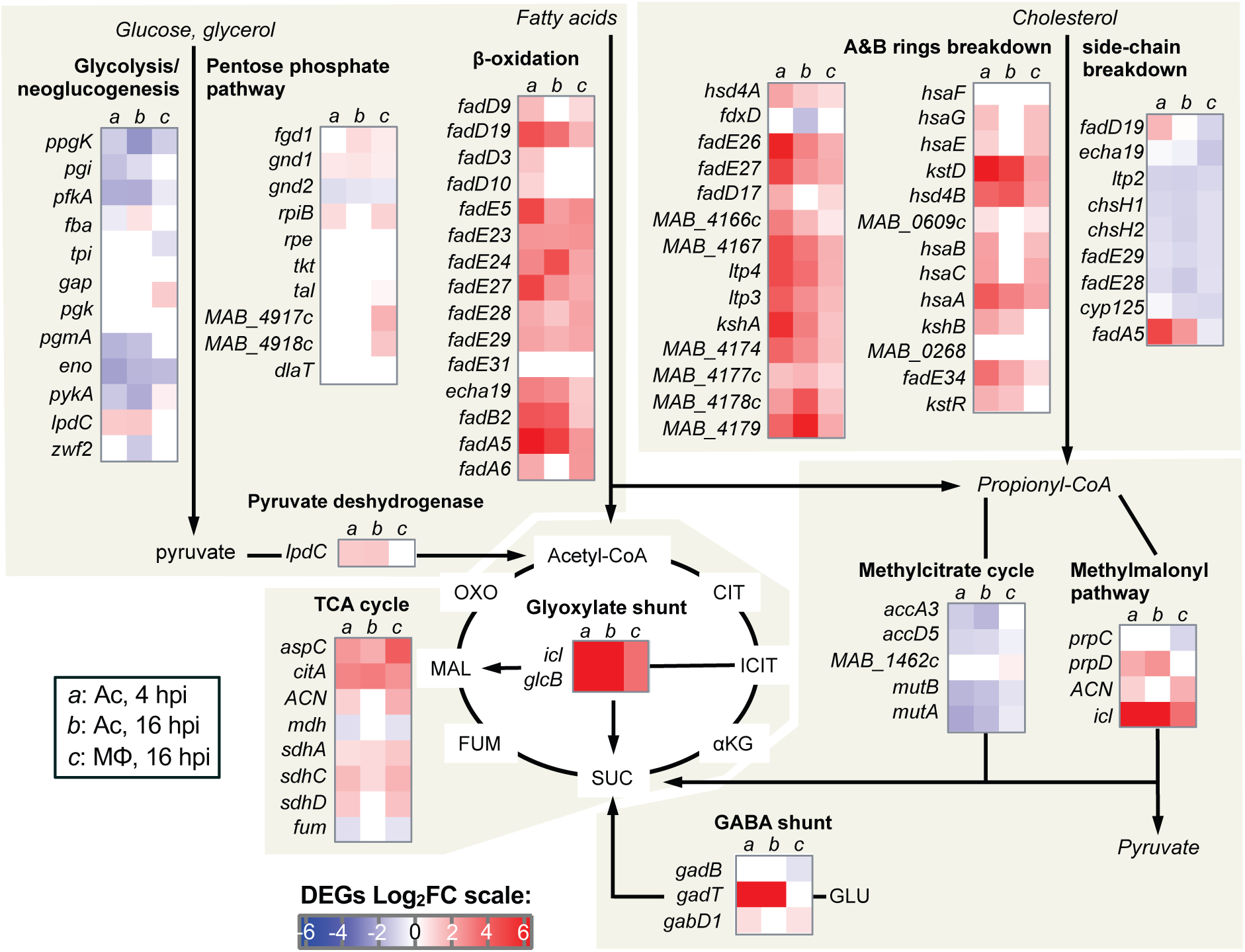
Intracellular *M. abscessus* relies on fatty acid and cholesterol catabolisM. DEGs FC of genes implicated in Central Carbon MetabolisM (CCM) is represented on a Heat Map ranging froM blue (DOWN DEGs) to red (UP DEGs). On this Heat Map both *M. abscessus* intra-amoebal (Ac) and intra-Macrophagic (Mφ) DEGs are depicted.

Furthermore, down-regulation of the mycolate operon (*MAB_2027*-*MAB_2039*) (**Table 1**), encompassing the β-ketoacyl-ACP synthases (KasA & KasB) and β-ketoacyl synthases (MAB_2031 & MAB_2029), as well as the malonyl-CoA acyl carrier protein transacylase (MCAT) homolog (MAB_2034), revealed that intracellular *M. abscessus* undergoes starvation as previously described (Jamet et al., 2015).

**Table 1:** Regulation of *M. abscessus* mycolate synthesis 882 operon in Ac and M⟴.

| **Mma gene** | **Encoded protein** | **Mabs gene** | **FC Ac**<br>**4 hpi** | **FC Ac**<br>**16 hpi** | **FC Mφ**<br>**16 hpi** |
| --- | --- | --- | --- | --- | --- |
| *MYCMA_RS13950* | Acyl carrier protein | *MAB_2027* | -3.28 | -2.91 | 0.00 |
| *MYCMA_RS13945* | 3-oxoacyl-ACP synthase (Kas B) | *MAB_2028* | -2.68 | -2.24 | 0.00 |
| *MYCMA_RS13940* | Beta-ketoacyl synthase | *MAB_2029* | -2.95 | -4.48 | 0.00 |
| *MYCMA_RS13935* | 3-oxoacyl-ACP synthase (Kas A) | *MAB_2030* | -3.80 | -3.15 | 0.00 |
| *MYCMA_RS13930* | Beta-ketoacyl synthase | *MAB_2031* | -2.92 | 0 | -1.60 |
| *MYCMA_RS13925* | 3-oxoacyl-ACP reductase | *MAB_2032* | -2.16 | 0 | -1.85 |
| *MYCMA_RS13920* | Thioesterase | *MAB_2033* | -2.09 | 0 | 0.00 |
| *MYCMA_RS13915* | Malonyl CoA-ACP transacylase | *MAB_2034* | -2.09 | 0 | 0.00 |
| *MYCMA_RS13910* | Acyltransferase | *MAB_2035* | 0 | 0 | 0.00 |
| *MYCMA_RS13905* | Membrane protein (MmpS) | *MAB_2036* | -1.16 | 0 | -1.06 |
| *MYCMA_RS13900* | Hypothetical protein (MmpL) | *MAB_2037* | -0.68 | -0.86 | 0.00 |
| *MYCMA_RS13895* | Transporter | *MAB_2038* | 0 | 0 | 0.00 |
| *MYCMA_RS13890* | Lipase (LipH) | *MAB_2039* | 0 | 0.86 | -0.72 |

### Regulation of putative virulence genes of M. abscessus *in Ac and Mφ*

We assessed the regulation of *M. abscessus* genes conserved in *M. tuberculosis* that are known to be induced and to contribute to the cellular microbicidal defenses of the tubercle bacillus in Mφ (Mukhopadhyay et al., 2012). The picture looked similar between Ac and Mφ with a few exceptions in the response to low O_2_/NO and in the low iron response (**Figure 4**). Transcriptional regulators such as *dosR*, *phoP* and *mtrA* were regulated in the opposite direction in Ac and Mφ, with *phoP* and *mtrA* only induced in Ac, and *dosR* exclusively induced in Mφ. Other genes known to contribute to the survival of the bacterium in response to oxidative stress (Sherman et al., 1999), comprising *ahpD*, *bcp*, *trxB1* and *2*, *trxC* genes (Schnappinger *et al.*, 2003), in addition to *ahpC*, were up-regulated in Ac and Mφ.

**Figure 4:**
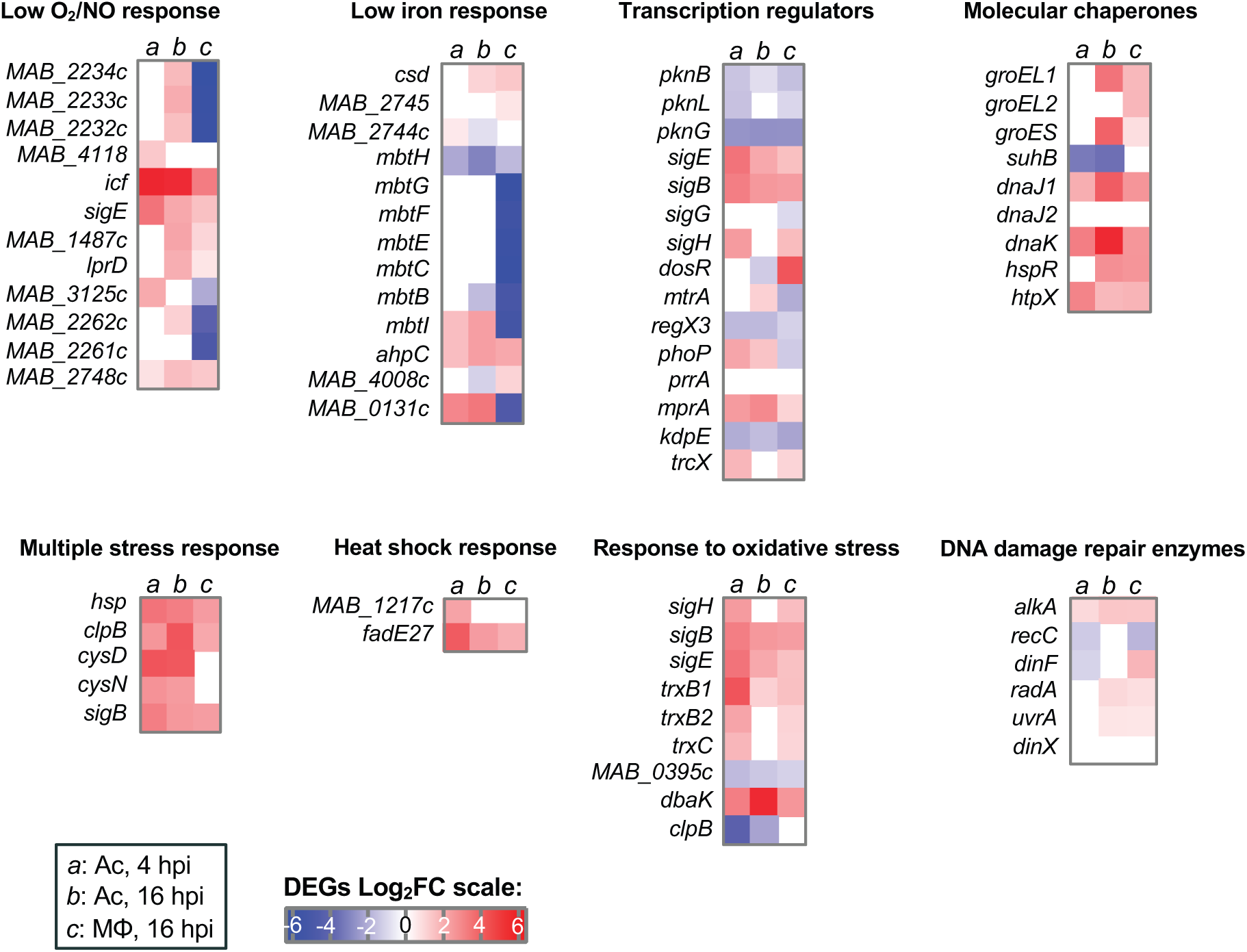
Regulation of genes required for pathogenic Mycobacteria survival *in vitro*. Regulation of *M. abscessus* genes conserved in *M.* tuberculosis, known to be induced and to contribute to cellular Microbicidal defenses of the tubercle bacillus in M (Mukhopadhyay et al., 2012) is represented on a Heat Map in a blue (repressed genes) to red (induced genes) color scale. On this Heat Map both *M. abscessus* intra-amoebal (Ac) and intra-Macrophagic (Mφ) DEGs are depicted and divided in categories: « broad transcription regulators » and genes implicated in the response to various intracellular stress (« Multiple stress response », « Heat Shock response », « Molecular chaperones », « DNA damage repair enzymes », « Response to low O_2_ / NO », « Low iron response », « Response to oxidative stress »).

Altogether, these analyses suggest that the induced sets of genes in Ac reflect the main adaptations to resistance to intracellular stress that were also shown to be induced in Mφ, hence suggeting that they constitute a repertoire of genes participating in *M. abscessus* virulence through intra-phagocyte survival.

### M. abscessus *highly-induced genes in amoeba*

The highest FC values of the *M. abscessus* in Ac 4 and 16hpi were chosen and then compared to the FC values obtained from the *M. abscessus* transcriptomes in Mφ and the *M. chelonae* transcriptome in Ac (**Sup. Figure 3**). This comparison highlighted 45 *M. abscessus* genes (**Sup. Table 2**). We also listed the most induced genes in Mφ (38 genes) (**Sup. Table 3**). From the comparison with the Ac transcriptomes, we constructed deletion mutants in 6 loci (ΔOP1 to ΔOP6) (**Table 3**). From the *M. abscessus* transcriptome in Mφ we constructed deletion mutants in 5 loci (ΔOP 7 to Δ OP11) among the most induced genes or in genes implicated in the adaptation to intracellular stress (**Table 3**).

ΔOP1 to ΔOP6 strains were evaluated for their intracellular Multiplication in Ac and in M (**Figure 5A**). All mutants were attenuated in Ac and Mφ, except one (ΔOP5), which multiplied More than the wild-type (wt) strain in M (**Figure 5A**). All mutated strains had similar growth *in vitro* to the wild type strain (**Sup. Figure 4A**). ΔOP2, 3, 4 and 6 were complemented and all strains recovered the wt phenotype except for the OP4 gene *MAB_2650* potentially encoding an MmpL (**Sup. Figure 4B**).

**Figure 5:**
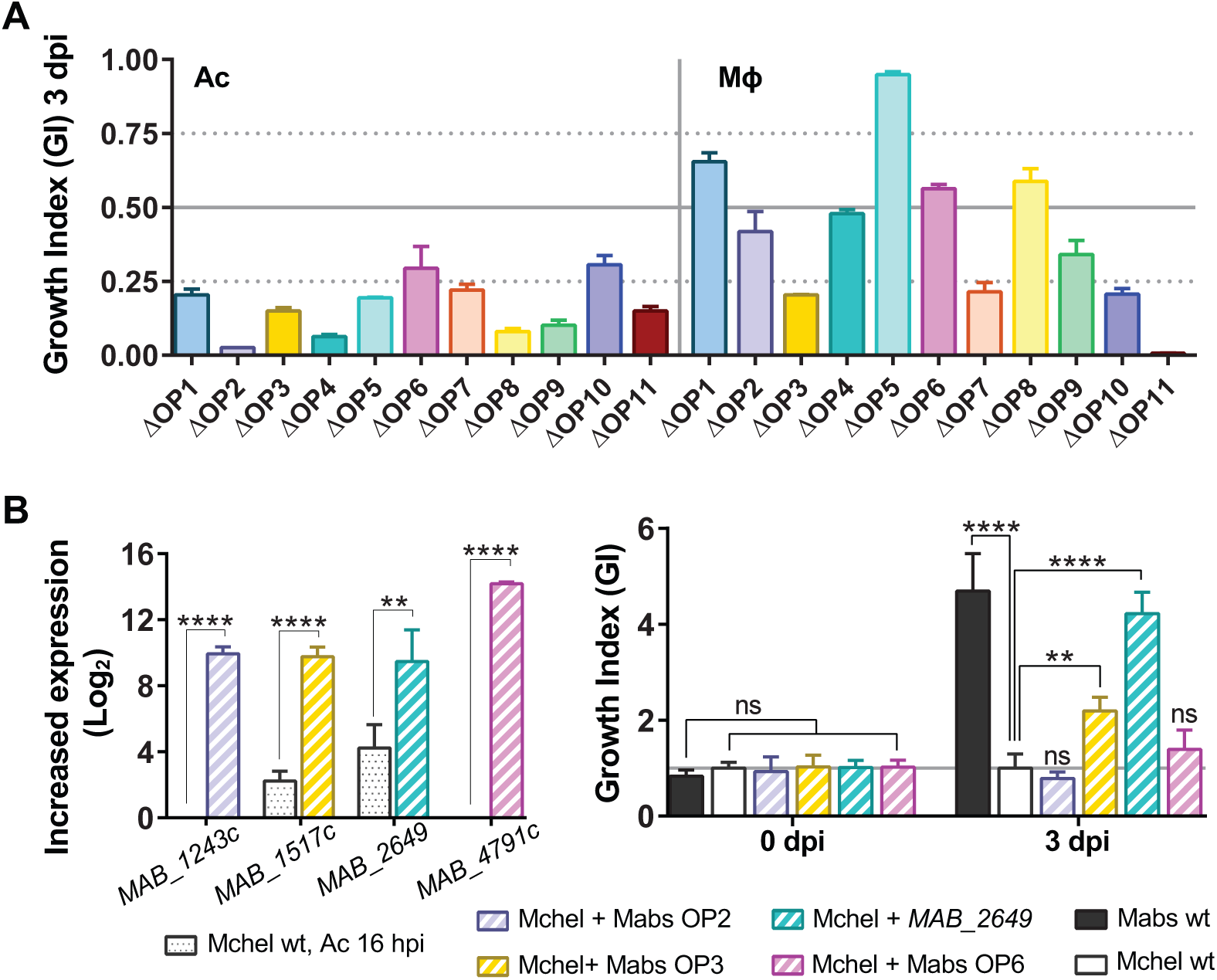
Comparative transcriptomic analyses allow identifying genes required for *M. abscessus* survival in amoebae and Macrophages. **A**. Intracellular survival of selected KO (∆OP) in *A. castellani* (Ac) and Macrophages (Mφ). **B**. Over-expression of 4 *M. abscessus* virulence genes (left panel) in *M. chelonae* and impacts on *M. chelonae* replication in Macrophages (right panel). FC values From *M. chelonae* transcriptome in Ac 16 hpi are compared to FC value Mid-log phase cultures of *M. chelonae* strains overexpressing *M. abscessus* genes (left panel). Cells were infected at 10 MOI and colony forming units (CFU) tests were performed 0 and 3 dpi. The relative growth of each strain as compared to *M. abscessus* wt (Growth Index, GI) is given. All experiments were repeated twice or more, in duplicates (A.) or triplicates (B). Statistical analyses were performed with GraphPad PRISM6. Histograms with error bars represent Means ± SD. Differences between means were analyzed by ANOVA and the Tukey post-test allowing Multiple comparisons to be performed. ns = non-significant. * *p*<0.05. ** *p*<0.01. *** *p*<0.001. **** *p*<0.0001.

These experiments confirmed the importance of these highly induced genes in Ac, in the intra-Mφ survival of *M. abscessus*. Several genes (OP2 and 6) are absent from *M. chelonae*, or present in the *M. chelonae* genome but at least four times less induced (OP3 and OP4) in Ac. We have analyzed their contribution towards intracellular survival by overexpressing *M. abscessus* OP2, 3, 6 and *MAB_2649* genes in *M. chelonae* (**Figure 5B, left panel**), which is unable to survive in Mφ (Sousa et al., 2015). Only the overexpression of *M. abscessus* OP3 and OP4 (*MAB_2649*) increased *M. chelonae* survival in Mφ (**Figure 5B, right panel**). By comparison, no increase in *M. chelonae* intracellular survival was observed when overexpressing OP2 and OP6 (**Figure 5B, right panel**).

### M. abscessus *highly-induced genes in Mφ*

The observed defect in intracellular survival was noticed for all mutants from OP7 to OP11 (**Figure 5A**), the ΔOP11 Mutant was particularly attenuated (GI<0.1). OP11 (*MAB_4532c*) KO strain tended to be eliminated by Mφ while the KO growth was similar to the WT growth *in vitro* (**Sup. Figure 4**). When complementing OP11 strain with *MAB_4532c*, we restored the wt phenotype (**Figure 6A**). *MAB_4532c* encodes an Eis N-acetyl transferase protein, according to a motif analysis (InterProScan 5). Of interest, *M. abscessus* contains two *eis* genes, named *eis1_MAB_* (*MAB_4124*) and *eis2_MAB_* (*MAB_4532c*) while *M. tuberculosis* possesses a single *eis_MTB_* gene (*Rv2416c*), *eis1_MAB_* (*MAB_4124*) being the closest homolog by Bidirectional Best Hit (BBH) search. No conservation of synteny was observed between the respective genomic regions in *M. abscessus* and *M. tuberculosis* (**Sup. Figure 5A**). In contrast, the *eis2_MAB_* locus shows some similarity and conservation with the *M. tuberculosis mmpL11* locus, with inverted and syntenic conservation of groups of genes (**Sup. Figure 5B**). The *eis_MTB_* locus was well-conserved in *M. abscessus* and corresponds to *MAB_1619*-*MAB_1627* and *MAB_1633*-*MAB_1637* regions (**Sup. Figure 5C**). Both *eis1_MAB_* and *eis2_MAB_* were found close to *mmpL* (brown arrows) and/or *mmpS* (orange arrow) genes (**Sup. Figure 5**). In the *M. abscessus eis2* locus, an *mmpL* gene (*MAB_4529*) was conserved in *M. tuberculosis* corresponding to *mmpL11* (**Sup. Figure 5B**). To assess the function of *M. abscessus eis* genes we performed transcomplementation of KO strains (*eis1_MAB_* and *eis2_MAB_*) with *M. tuberculosis eis* (*eis_MTB_*). Of interest, and unlike *eis2_MAB_*, *eis1_MAB_* was suppressed inside M (**Sup. Figure 6**) and less impaired in its intracellular survival (**Figures 6A-B**). Complementation of *eis1_MAB_* and*eis2_MAB_* with *eis1_MAB_* or *eis2_MAB_* respectively allowed the recovery of the intracellular survival for both mutants (**Figures 6A-B**). However, Complementation of both mutants with the *eis_MTB_* gene allowed only partial restoration of the intracellular replicative phenotype for the *eis2_MAB_* Mutant, but no restoration was observed for *eis1_MAB_* mutant (**Figure 6A-B**). similar behaviors regarding apoptosis, necrosis, autophagy and phagosomal acidification were observed when comparing the wt *M. abscessus* strain with the *eis2_MAB_* mutant (**Sup. Figure 7**). However, two Major differences were observed. First, infection of Mφ with the *eis2_MAB_* strain (at a MOI of 50) was associated with higher production of ROS by the cells and loss of *eis2_MAB_* also sensitized *M. abscessus* to ROS and notably to H_2_O_2_ (**Figures 6C-D**). Secondly, the *eis2_MAB_* mutant was unable to damage the phagosomal Membrane and to provoke phagosome-cytosol contact as compared to the wt and complemented *M. abscessus* strains (**Figure 6E**).

**Figure 6:**
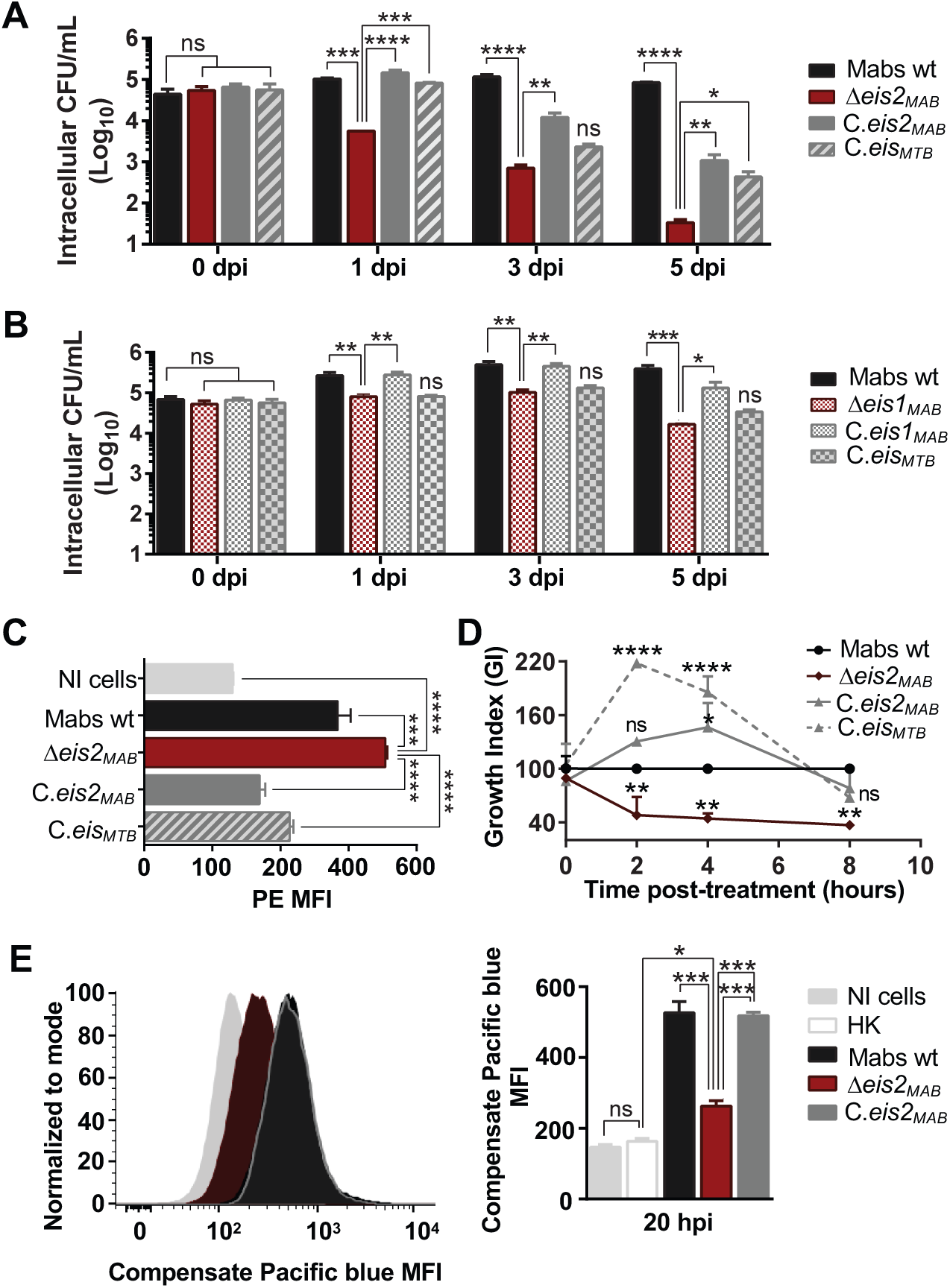
*M. abscessus eis2* gene is essential for survival in Macrophages and shares functions with *M. tuberculosis eis* conversely to *M. abscessus eis1*. **A**. Intracellular survival of *M. abscessus eis2* KO strain (∆*eis2_MAB_*) and Complementation in macrophages (Mφ). **B**. Intracellular survival of *M. abscessus eis1* KO strain (∆*eis1_MAB_*) and Complementation in Mφ. macrophages were infected at 10 MOI and colony forming unit (CFU) tests were performed at several times post-infection (0, 1, 3 and 5 dpi). **C**. Control of ROS production by *M. abscessus* Eis2. ROS production by Mφ was assessed by flow cytometry with the Mitosox Red kit, 15 Min post-infection at 50 MOI. **D**. Sensitivity of *M. abscessus eis2* KO strain to hydrogen peroxide (H_2_O_2_). Sensitivity to H_2_O_2_ was assessed by incubated bacterial cultures with H_2_O_2_ 20 µM during 8 h. The amount of survival cells was determined by performing CFU tests at several hours post-infection (2, 4, 8 hpi). E. *Control of phagosomal rupture by M. abscessus Eis2.* phagosomal rupture was assessed by performing a Fluorescence energy transfer (FRET) analysis as previously described (Simeone *et al.*, 2015). Results are depicted as signal overlays per group with 1,000,000 events per condition acquired in not infected cells (NI cells), Heat killed *M. abscessus* (HK), wild-type *M. abscessus* (Mabs wt), KO strains (Δeis2_MAB_), KO strains complemented with eis2_MAB_ (C.eis2_MAB_). All experiments were repeated twice or more in triplicates. Statistical analyses were performed with GraphPad PRISM6. Histograms with error bars represent means ± SD. Differences between means were analyzed by ANOVA and the Tukey post-test allowing multiple comparisons to be performed. ns = non-significant. * *p*<0.05. ** *p*<0.01. *** *p*<0.001. **** *p*<0.0001.

## Discussion

The main objective of this work was to understand the genetic and molecular basis for the ability of *M. abscessus* to withstand and survive in eukaryotic phagocytic cells. Identification of genes strongly induced after infection of Ac allowed us to reveal those genes that play key roles in Mφ intracellular survival. Conversely, identification of genes strongly induced after infection of Mφ allowed us to reveal those essential to Ac intracellular survival with the exception of one mutant, OP5 (*MAB_4663*) encoding a protein of unknown function, that showed enhanced growth in Mφ. KO mutants constructed on the basis of the results from transcriptomic analyses allowed the identification of genes required for bacterial survival in phagocytes. These results are complementary to a previous *Tn M. abscessus* library viability screen in Ac, permitting the identification of two other intracellular virulence factors, namely the type VII secretion system ESX4 and the lipid transport protein MmpL8_MAB_ (Dubois et al., 2018b; Laencina et al., 2018). The intracellular defects of strains that were deleted for genes highly induced in amoeba (induced at least four times more compared to the intramacrophagic transcriptome of *M. abscessus* and to the intra-amoebal transcriptome of *M. chelonae* (OP1 to OP6)), suggest that the transcriptomic changes observed following a co-culture in amoebae reflected the response of *M*. *abscessus* in Mφ. In addition, co-culturing *M. abscessus* in amoebae enhances the virulence of *M. abscessus* even *in vivo* through the induction of the PLC virulence gene (Bakala N’Goma *et al.*, 2015). These transcriptomic analyses highlight that exposure to the amoebal intracellular environment potentiates *M*. *abscessus* virulence, increasing the resistance of the bacteria for future encounters with phagocytic cells that form part of the innate defense of the host.

Most of the 6 loci (OP1 to OP6) studied encode for hypothetical proteins, with the exception of the *MAB_1517c* gene that encodes a probable O-methyltransferase (OP3), and the *MAB_2649*and *MAB_2650* genes encoding an MmpS and an MmpL, respectively (OP4) (**Table 2**) (Viljoen et al., 2017). At the *M. abscessus* OP2 locus, that mainly comprises genes of unknown function, we used a motif analysis to identify an ABC transporter, FecCD/TroCD-like in the MAB_1243c protein and an alkaline shock protein Asp23 in the MAB_1247c protein. A Motif analysis performed on *M. abscessus* OP6 gene shows that *MAB_4791c* encodes a protein implicated in the twin-arginine translocation pathway (see below). Notably, the over-induction of *MAB_2649* and *MAB_1517c* in *M. chelonae* favors its replication in Mφ, suggesting that the high induction of these two genes in amoeba may trigger *M. abscessus* virulence.

**Table 2:**
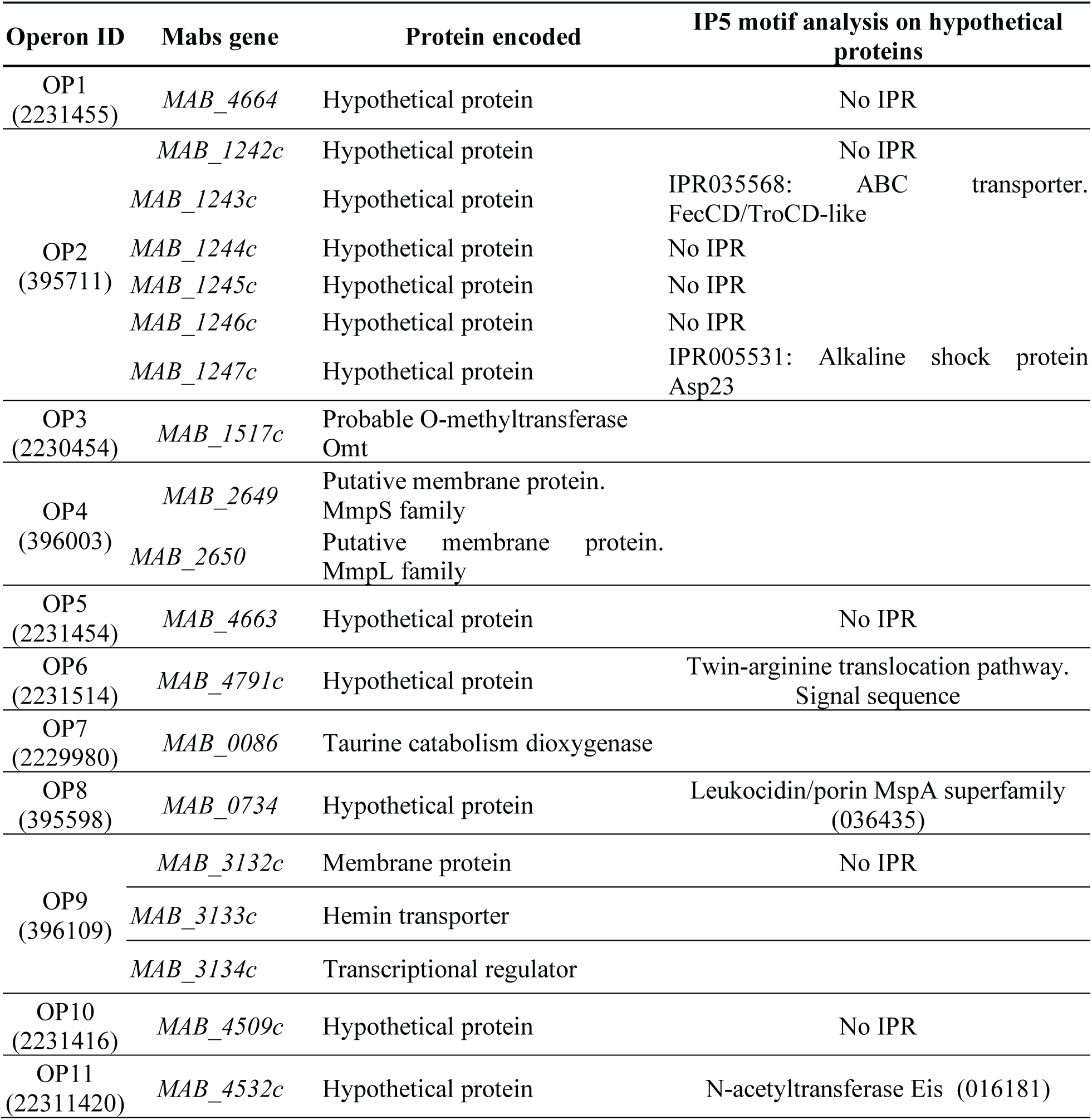
Deleted operons in *M. abscessus* ATCC19977.

A striking finding of our work is the essential role of the *eis2_Mab_* gene in early resistance to the Microbicidal action of Mφ, via phagosomal Membrane damage and cytosol contact, that allows the intracellular survival of *M. abscessus*. Although *M. abscessus* possesses two *Eis* genes, there is no redundancy in their respective functions; the *eis1_Mab_* mutant presented a similar behavior to the wt strain in Mφ, with only the loss of a log_10_ CFU at 5 dpi, compared to the quasi-total clearance of the *eis2_Mab_* mutant in Mφ. Despite higher genomic identity between *eis1_MAB_* and *eis_MTB_*, the restoration of the phenotype when complemented with *eis_MTB_* was observed only for the *eis2_MAB_* mutant, demonstrating the similar role of *eis2_MAB_* to what is described for *eis_MTB_* in virulence. However the deletion of this gene in *M. abscessus* is more deleterious for the bacterium in Mφ compared to the deletion of *eis_MTB,_* (Wei *et al.*, 2000; Wu *et al.*, 2009). E*is_MTB_* gene has been described as being important for *M. tuberculosis* survival inside Mφ by controlling host cell apoptosis, autophagy, ROS production and innate immune defenses (Shin et al., 2010). As also observed in *M. tuberculosis* (Shin et al., 2010), increasing the MOI (to 50) revealed further differences between *eis2_MAB_* KO and wt strains with regard to resistance to oxidized derivatives; however, this bacterial-load effect has not yet been observed for cell death mechanisms.

One of the peculiarities of the locus *eis2_MAB_* is to possess similarities with a genomic region of *M. tuberculosis*, within which is found the gene *mmpL11*. The potential counterpart in *M. abscessus* would be *MAB_4529* (**Sup. Figure 5**). Most of MmpL are lipid transporters implicated in cell physiology and virulence (Chalut, 2016; Viljoen et al., 2017). *M. abscessus* has 27 MmpLs, twice as much as *M. tuberculosis* (Viljoen et al., 2017). In *M. tuberculosis*, MmpL11 is implicated in heme iron acquisition (Owens et al., 2013) and transport of mycolic acid wax ester and long-chain triacylglycerols (Wright et al., 2017). Three genes conserved in the *M. abscessus eis2* locus encode for proteins belonging to lipid transport and metabolism pathways (COG I), which suggests, together with the conservation seen with the *M. tuberculosis mmpL11* locus, that the *M. abscessus eis2* locus might also participate in cell wall biogenesis (Yamaryo-Botte et al., 2015).

Transcriptomic analysis revealed that nine *M. abscessus* genes, whose orthologues in *M*. *tuberculosis* contribute to virulence, were highly induced during infection of Mφ (**Sup.Table 2**). Among their gene products, WhiB7 and DevR-DevS are implicated in stress sensing (Kumar et al., 2011). WhiB7, a Fe-S cluster protein, was shown to be induced in response to perturbation in amino-acid metabolism, under reducing intracellular state, iron depletion and increased temperatures (Geiman et al., 2006). The 20-fold increase of *M. abscessus whiB7* in Mφ suggests that *M. abscessus* may undergo similar stresses in Mφ. The DevR response regulator of the histidine kinase DevS was also highly up-regulated. In *M. tuberculosis*, the *devR-devS* two-component system (also known as the DosR system) is activated in response to hypoxia (Sherman et al., 2001). Likewise, *M. abscessus MAB_2562c,* the orthologue of *Rv0081,* was induced 10-fold in Mφ. A putative orthologue (*MAB_1409c*) of the dormancy response gene *Rv1258c* was also strongly induced in intra-macrophagic *M. abscessu*s. The conserved alpha-ketoglutarate-dependent dioxygenase AlkB-encoding gene is thought to be involved in fatty acid metabolism, or in protection against DNA methylation. The *aspC* gene was induced 8-fold; AspC mediates nitrogen transfer from aspartate to glutamate, which in turn, together with glutamine, provides nitrogen to most of the biosynthesis pathways. This is thought to be essential in *M. tuberculosis* (Sassetti et al., 2003), while aspartate is required for mycobacterial virulence (Gouzy et al., 2013). *M. abscessus katA* gene, which is conserved in *M. aviuM* and *Listeria monocytogenes*, is a catalase that degrades H_2_O_2_ into water and oxygen in a single reaction. Such a reaction, enabling resistance to oxidative metabolites, may be an important mechanism of bacillary survival within the host phagocyte (Manca et al., 1999). *M. abscessus eaMA (MAB_0677c)*, which is thought to encode a drug/metabolite transporter, was induced in Mφ. Two additional genes (*MAB_3762* and *MAB_*3180) encoding proteins with an EaMA domain were also highly induced. Finally, at the molecular function level, it appears that six of the most highly induced genes in *M. abscessus* in Mφ encode acyl or N-acetyl transferase proteins playing a role in post-translational modifications.

The *M. abscessus* transcriptomes’ comparison in Ac or Mφ allowed differences in metabolic adaptations to be highlighted. In Mφ, *M. abscessus* enters a slow replicative stage, and activates the detoxification and protein secretion pathways. By comparison, in amoebae *M. abscessus* switches on protein synthesis, lipid transport and metabolism, transcription of genes involved in post-translational modifications (PTM), protein turnover and chaperones (COG O), reflecting a more active and replicative behavior as compared to a more persistent state in M. Actually, cell wall biogenesis including peptidoglycan and glycosaminoglycan biosynthetic processes were down-regulated in Mφ. Similarly, *mtrA*, *phoP* and *devR* were differently regulated, with only *devR* up-regulated in Mφ, confirming the switch towards a slow growth stage for *M. abscessus* in Mφ.

Over-representation of the COG O (post-translational modification, protein turnover, molecular chaperone) category in *M. abscessus* infecting Ac indicates that *M. abscessus* may alter cellular processes during its interactions with host cells via PTM, as described in various pathogens (Ribet and Cossart, 2010; Müller *et al.*, 2010; Parra *et al.*, 2017). Protein turnover does not only help in clearing of old proteins but also aids a fast adaptation to nutrient poor environments (Goldberg and St. John, 1976). Molecular chaperones help pathogens override unfavorable conditions found in the host such as heat shock, oxidative and acid stresses (Neckers and Tatu, 2008). They also contribute to the inhibition of lysosomal fusion and favor bacterial growth (Neckers and Tatu, 2008). Molecular chaperones may therefore form a first line of defense and help consolidate pathogen virulence. Thus, over-representation of the COG O inside amoebae might reflect specific intracellular cues the mycobacterium faced from the early time points post infection.

In Ac, the most enriched GO is adenine salvage (GO:0006168) (**Figure 2**). This GO represents any process that generates adenine from derivatives without any *de novo* synthesis. Mycobacteria are able to limit the synthesis of this high energy demanding nucleotide (Ducati et al., 2011). Mycobacteria are also capable of scavenging free nitrogenous bases From the medium (Ducati et al., 2011). Under conditions of low energy availability or rapid multiplication, the salvage pathway may then be the main source of maintaining the nucleotide pool (Ducati et al., 2011).

Sulfur metabolism (GO:0000103), hydrogen sulfide (H_2_S) biosynthetic pathway (GO:0070814) and detoxification via Fe-S cluster assembly proteins (GO:0016226) in addition to polyamine transport (GO:0015846), were also enriched by *M. abscessus* in Ac. In its reduced form, sulfur is used in the biosynthesis of the amino acid cysteine that is one of the prime targets for reactive nitrogen intermediates (Rhee et al., 2005). Those pathways might play a key role in *M. abscessus* survival in phagocytic cells, since genes involved in the metabolism of sulfur have consistently been identified as up-regulated in conditions that Mimic the intra-macrophagic environment and during Mφ infection for *M. tuberculosis* (Schnappinger *et al.*, 2003). As for polyamines (cadaverine, putrescine and spermidine), they are known to have pleiotropic effects on cells via: their interaction with nucleic acids; a role in bacterial virulence by allowing mycobacterial escape from the phagolysosome; toxin activity or protection from oxidative and acid stress has also been demonstrated (Shah and Swiatlo, 2008).

In Mφ, glycerol ether metabolic process, MEP pathway and L-proline biosynthetic processes were the most enriched. Glycerol ether metabolic process corresponds to glycerophospholipids-seminolipids-plasmecholine metabolism and cellular amide biosynthetic processes. The MEP pathway is required for isoprenoid precursor biosynthesis (Rohmer et al., 1993). A wide variety of Monoterpenes and diterpenes belong to isoprenoid classes which function as toxins, growth inhibitors, or other secondary metabolites (Gershenzon and Dudareva, 2007). Finally, proline has been reported as an important factor in the adaptation of mycobacteria to slow growth rate and hypoxia (Berney *et al.*, 2012). It is believed that the proline-utilization pathway protects mycobacterial cells by detoxifying methylglyoxal, a by-product of endogenous glycerol metabolism (Berney *et al.*, 2012) that can damage DNA and proteins within cells. Up-regulation of base-excision repair suggests that intracellular mycobacteria undergo DNA damage. Protein folding was also enriched, as well as the type II secretion system, which was enriched by more than two-fold. This secretion system promotes the specific transport of folded periplasmic proteins across a dedicated channel in the outer membrane, and it facilitates both Sec and Tat pathways to secrete proteins into the periplasm. Potential roles for SecA1 and SecA2 in *M. tuberculosis* dormancy has been reported while the Tat pathway was shown to contribute to virulence in *Legionella pneumophila* for instance, by aiding secretion of Phospholipase C (Rossier and Cianciotto, 2005), a virulence factor conserved in *M. abscessus* (Ripoll *et al.*, 2009).

Both Ac and Mφ were sensed as a stressful environment by *M. abscessus,* evidenced by the up-regulation of genes known to be involved in multiple stress responses. Induction of low O_2_ and low NO response genes confirm that hypoxic environments are encountered by *M. abscessus* both in Ac and Mφ.

In conclusion, our findings confirm that the amoeba-induced genes play a role in potentiating the subsequent survival of *M. abscessus* in Mφ. Both environments have commonalities, in terms of metabolic switches, especially to withstand the host response. It is through this preparation during its intra-amoebic life that *M. abscessus* is able to withstand the noxious M environment, especially thanks to several genes whose role has been confirmed during this work. The multiple leads opened during this work must now be followed to complete this viewpoint of synergistic potentiation of virulence conferred by the amoeba to *M. abscessus*, including the ultimate mechanisms of manipulation of the host’s defense systems as seen with other intracellular pathogens.

## MATERIALS AND METHODS

### Bacterial strains, plasmids and growth conditions

A clinical isolate of *M. abscessus* subspecies *Massiliense s*Mooth variant (43S) and *M. chelonae* type strain CCUG 47445 were used for the RNAseq experiments. Gene deletions were performed with CIP 104536T type smooth strain of *M. abscessus* subspecies *abscessus*. Both *M. abscessus* CIP strain and *M. chelonae* type strain were used to perform *in vitro* survival and Complementation tests while gene deletions were performed with *M. abscessus* CIP 104536T strain. *M. abscessus* and *M. chelonae* strains were routinely grown aerobically at 37 °C and 32°C respectively, in Middlebrook 7H9 medium (Sigma-Aldrich) supplemented with 0.2% glycerol, 1% glucose, and 250 Mg/L kanamycin (Thermo Fisher Scientific) when necessary, with 25 Mg/L zeocin (Thermo Fisher Scientific) for the knockout strains, and with 25 Mg/L zeocin plus 250 Mg/L hygromycin (Thermo Fisher Scientific) for complemented strains. *A. castellanii* (ATCC 30010) was grown at rooM temperature without CO_2_ in peptone yeast extract glucose (PYG) broth for the amplification of the strain. J774.2 cell line was grown and used as described (Le Moigne et al., 2016; Roux et al., 2016).

### Gene deletion and Complementation

Deletion of genes was performed using the recombineering system as described previously (Medjahed and Singh, 2010; Bakala N’Goma *et al.*, 2015). Growth of the KO strains was checked by Measuring the optical density of bacterial cultures in 7H9 medium supplemented with glycerol 0.1%. Complementation was performed after amplifying and cloning genes into the integrative plasmid pMVH361 as described (Bakala N’Goma *et al.*, 2015).

### RNA isolation and RNA sequencing

Approximately 10^7^ cells were infected in 50 ML tubes, with low agitation, without CO_2_. amoebae were infected at 100 MOI at 32°C. M were infected at 50 MOI at 37°C. Cells were washed 3 times after 1 hour of infection and resuspended in medium supplemented with amikacin 250 µg/ML and incubated for 1 hour to eliminate extracellular bacteria. Three additional washed were performed and cells were resuspended in medium supplemented with amikacin 50 µg/ML for the rest of the infection. amoebal cells were harvested 4 h and 16 hpi for intracellular *M. abscessus* RNA isolation 4h and 16 hpi and 16hpi for intracellular *M. chelonae* RNA isolation. M were harvested for intracellular *M. abscessus* RNA isolation 16H post-infection. RNA isolation was performed as described (Dubois et al., 2018a). Briefly, cells were lysed with a cold solution of GuanidiuM thiocyanate (GTC), N-Lauryl-sarcosine, SodiuM citrate +/- Tween 80 plus-Mercaptoethanol. The lysates containing intracellular bacteria were collected, centrifuged and RNA was isolated from the bacterial pellets with TRIzol. The lysates were then transferred into 2 ML screw tubes containing zirconiuM beads and were conserved at −80°C for at least 1 day to allow inactivation of RNAses and cells dissolution. Bacteria cells were disrupted with a bead beater by performing to round at 6.500 rpM for 25 seconds, followed by one round at 6.500 rpM for 20 seconds. Two hundred µL of chloroforme isoamyl was added and tubes were iMMediately Mixed for 10 seconds. The Mixture was centrifugated at 13.000 rpM for 15 Minutes at 4°C. The RNA present in the upper phase was transferred to a fresh tube and precipitated by adding 0.8 volume of isopropanol. Tubes were inverted twice to allow precipitation and kept at −20°C for at least 2 hours. The precipitated RNA was then pelleted by centrifugation at 13.000 rpM for 30 Min. at 4°C. The pellet was washed with ethanol (70%) and centrifuged at 13.000 rpM for 10 Min at 4°C. The washed pellet was air-dried, re-suspended in RNase-free water and stored at −80°C until cDNA library construction.

Control RNA was isolated from bacteria cells grown in amoeba or M co-culture medium (Mowat and DMEM supplemented with 10% Fetal Bovine SeruM respectively). Biological replicates were prepared to allow statistical comparisons of infected and non-infected samples.

### RNA treatments prior to library preparation and library preparation

RNA samples were treated with DNases (AMBION) to remove DNA contaminants, purified with the RNA MEGAclear kit (Thermofisher), and depleted of ribosomal RNA with the riboZero kit (Illumina). RNA (total, depleted, purified) is checked on the Bioanalyser system (Agilent) for its quality and integrity. cDNA libraries were prepared with samples displaying a RIN above 7. RNA concentrations were Measured using the nanodrop spectrophotometer (Thermo Scientific) and the Qubit fluorometer (Invitrogen). Libraries were prepared with the TruSeq Stranded RNA LT prep kit cDNA synthesis, set A (Illumina) which consists in: (1) RNA fragmentation, (2) 1st strand cDNA synthesis (Reverse transcriptase and randoM primers), (3) 2d strand cDNA synthesis (removal of the RNA template and synthesis of a new strand with dUTP), (4) no end repair step, (5) adenylation of 3’ ends, (6) ligation of adapters and (7) enrichment of DNA fragments. Libraries are checked for concentration and quality on DNA chips with the Bioanalyzer Agilent. more precise and accurate quantification is performed with sensitive fluorescent-based quantitation assays (“Quant-It” assays kit and QuBit fluorometer, Invitrogen).

### NGS sequencing and data analysis

Sequencing and statistical analysis were performed in the transcriptome and Epigenome platform (PF2) of Pasteur Institut, Paris, France. The cDNA libraries were sequenced on Illumina HiSeq 2500 system by performing an SRM run (SR: Single Read, PE: Paired-end Reads, M: multiplexed samples) of 51 cycles with 7 index bases read. The quality of the sequencing was assessed with the external FastQC program (https://www.bioinformatics.babrahaM.ac.uk/projects/fastqc/). After the triMMing of adapter sequences and low-quality reads with cutadapt version 1.11, reads were aligned with RefSeq assemblies, either *M. abscessus* subsp. *Massiliense* strain GO06 assembly (GCF_000277775.2) or *M. chelonae* CCUG 47445 assembly (GCF_001632805.1), using the Bowtie software version 0.12.7 (http://bowtie-bio.sourceforge.net/index.shtml) with defaults parameters. Genes were counted using featureCounts 1.4.6-p3 from Subreads package (parameters:-g gene-t ID-s 1). Differential analysis of gene expression was performed using the R software (version 3.3.1) and the Bioconductor packages DESeq2 (version 1.12.3) (Love et al., 2014) using the default parameters and statistical tests for differential expression were performed applying the independent filtering algorithM. A generalized linear Model was set in order to test for the differential expression between the biological conditions. For each pairwise comparison, raw *p*-values were adjusted for multiple testing according to the Benjamini and Hochberg (BH) procedure (Benjamini and Hochberg, 1995) and genes with an adjusted *p*-value lower than 0.05 were considered differentially expressed. Equivalent to *M. abscessus* subsp. *abscessus* CIP strain of *M. abscessus* 43S genes and *M. chelonae* genes were determined by Birectionnal Best Hit search using the Opscan software (http://wwwabi.snv.jussieu.fr/public/opscan/). Differentially expressed genes assignment to COGs was performed using the COG automatic Classification from the MicroScope database (Vallenet et al., 2009). The percentage assignments were compared by performing Fisher’s exact tests. GOE analyses were performed with the R software topGO package (Bioconductor) (Alexa *et al.* 2018).

### Quantitative real-time PCR

qRT-PCR were performed with a CFX96 thermal cycler (Bio-Rad). Controls without reverse transcriptase were done on each RNA sample to rule out DNA contamination. The sigA gene was used as an internal control (Bakala N’Goma *et al.*, 2015). Each qRT-PCR was performed with three biological replicates.

### In vitro survival assays

Survival of strains in amoebae and M were performed as previously described (Dubois et al., 2018b). Survival tests of KO strains were performed in duplicates three times. confirmation of attenuated phenotypes and Complementation tests were performed in triplicates three times.

### Phagosome acidification and phagosomal escape FRET assays Phagosome acidification

Phagosome acidification and phagosomal escape FRET assays and phagosomal escape FRET assays were conducted in THP-1 cells as previously described (Roux et al., 2016; Simeone et al., 2015).

### Cell death, autophagy and ROS production assays

Macrophage death following infection with *M. abscessus* was assessed with the Dead Cell Apoptosis Kit with Annexin V FITC and PI for flow cytometry (ThermoFischer). Autophagy was assessed with the Premo™ Autophagy TandeM Sensor RFP-GFP-LC3B Kit (ThermoFisher). ROS production by macrophages was Measured with the MitoSOX Red kit (ThermoFisher).

Infections were performed as previously described (Dubois et al., 2018b), at 50 MOI, except in the ROS production assay for which the cells were infected 15 Min only.

### Bacterial sensitivity to H_2_O_2_

Sensitivity to H_2_O_2_ was assessed by culturing the bacteria in 7H9 medium suuplemented with glycerol 0,1% and H_2_0_2_ 3% (Laboratoires Gillbert) (20 μM). CFU tests were performed at different times post-treatment (2h, 4h, 8h) to determine the number of viable bacteria compared to the wt strain (Growth Index).

## ACKNOWLEDGEMENT

We thank B. G. Marshall (Southampton University) and S. Gordon (University College, Dublin) for their careful reading of the Manuscript and for giving valuable coMMents.V.D. was supported by French Cystic Fibrosis Patients Association Vaincre la Mucoviscidose Grant RF20150501377.

## REFERENCES

Adékambi T, Ben Salah S, Khlif M, Raoult D, Drancourt M. 2006. Survival of environmental mycobacteria in Acanthamoeba polyphaga. Appl Environ Microbiol 72:5974–5981. doi:10.1128/AEM.03075-05

Ahmed N, Saini V, Raghuvanshi S, Khurana JP, Tyagi AK, Tyagi AK, Hasnain SE. 2007. Molecular Analysis of a Leprosy Immunotherapeutic Bacillus Provides Insights into Mycobacterium Evolution. PLoS One 2:e968. doi:10.1371/journal.pone.0000968

Angenent LT, Kelley ST, St Amand A, Pace NR, Hernandez MT. 2005. Molecular identification of potential pathogens in water and air of a hospital therapy pool. Proc Natl Acad Sci U S A 102:4860–5. doi:10.1073/pnas.0501235102

Bakala N’Goma JC, Le Moigne V, Soismier N, Laencina L, Le Chevalier F, Roux A-L, Poncin I, Serveau-Avesque C, Rottman M, Gaillard J-L, Etienne G, Brosch R, Herrmann J-L, Canaan S, Girard-Misguich F. 2015. Mycobacterium abscessus phospholipase C expression is induced during coculture within amoebae and enhances M. abscessus virulence in mice. Infect Immun 83:780–791. doi:10.1128/IAI.02032-14

Barker J, Brown MRW. 1994. Trojan Horses of the microbial world: protozoa and the survival of bacterial pathogens in the environment. Microbiology 140:1253–1259. doi:10.1099/00221287-140-6-1253

Ben Salah I, Adékambi T, Drancourt M. 2009. Mycobacterium phocaicum in therapy pool water. Int J Hyg Environ Health 212:439–444. doi:10.1016/j.ijheh.2008.10.002

Benjamini, Hochberg. 1995. Controlling the False Discovery Rate: A Practical and Powerful Approach to Multiple Testing. J R Stat Soc Ser B 57:289–300.

Berney M, Weimar MR, Heikal A, Cook GM. 2012a. Regulation of proline metabolism in mycobacteria and its role in carbon metabolism under hypoxia. Mol Microbiol 84:664– 81. doi:10.1111/j.1365-2958.2012.08053.x

Berney M, Weimar MR, Heikal A, Cook GM. 2012b. Regulation of proline metabolism in mycobacteria and its role in carbon metabolism under hypoxia. Mol Microbiol 84:664– 681. doi:10.1111/j.1365-2958.2012.08053.x

Chalut C. 2016. MmpL transporter-mediated export of cell-wall associated lipids and siderophores in mycobacteria. Tuberculosis (Edinb) 100:32–45. doi:10.1016/j.tube.2016.06.004

Cirillo JD, Falkow S, Tompkins LS, Bermudez LE. 1997. Interaction of Mycobacterium avium with environmental amoebae enhances virulence. Infect Immun 65:3759–67.

Dubois V, Laencina L, Bories A, Le Moigne V, Pawlik A, Herrmann J-L, Girard-Misguich F. 2018a. Identification of Virulence Markers of Mycobacterium abscessus for Intracellular Replication in Phagocytes. J Vis Exp. doi:10.3791/57766

Dubois V, Viljoen A, Laencina L, Le Moigne V, Bernut A, Dubar F, Blaise M, Gaillard J-L, Guérardel Y, Kremer L, Herrmann J-L, Girard-Misguich F. 2018b. MmpL8MAB controls Mycobacterium abscessus virulence and production of a previously unknown glycolipid family. Proc Natl Acad Sci U S A 201812984. doi:10.1073/pnas.1812984115

Ducati RG, Breda A, Basso LA, Santos DS. 2011. Purine Salvage Pathway in Mycobacterium tuberculosis. Curr Med Chem 18:1258–75.

Eddyani M, De Jonckheere JF, Durnez L, Suykerbuyk P, Leirs H, Portaels F. 2008. Occurrence of free-living amoebae in communities of low and high endemicity for Buruli ulcer in southern Benin. Appl Environ Microbiol 74:6547–53. doi:10.1128/AEM.01066-08

Falkinham JO. 2009. Surrounded by mycobacteria: nontuberculous mycobacteria in the human environment. J Appl Microbiol 107:356–67. doi:10.1111/j.1365-2672.2009.04161.x

Geiman DE, Raghunand TR, Agarwal N, Bishai WR. 2006. Differential Gene Expression in Response to Exposure to Antimycobacterial Agents and Other Stress Conditions among Seven Mycobacterium tuberculosis whiB-Like Genes. Antimicrob Agents Chemother 50:2836–2841. doi:10.1128/AAC.00295-06

Gershenzon J, Dudareva N. 2007. The function of terpene natural products in the natural world. Nat Chem Biol 3:408–414. doi:10.1038/nchembio.2007.5

Goldberg AL, St. John AC. 1976. Intracellular Protein Degradation in Mammalian and Bacterial Cells: Part 2. Annu Rev Biochem 45:747–804. doi:10.1146/annurev.bi.45.070176.003531

Gomila M, Ramirez A, Gasco J, Lalucat J. 2008. Mycobacterium llatzerense sp. nov., a facultatively autotrophic, hydrogen-oxidizing bacterium isolated from haemodialysis water. Int J Syst Evol Microbiol 58:2769–2773. doi:10.1099/ijs.0.65857-0

Gouzy A, Poquet Y, Neyrolles O. 2013. A central role for aspartate in Mycobacterium tuberculosis physiology and virulence. Front Cell Infect Microbiol 3:68. doi:10.3389/fcimb.2013.00068

Greub G, Raoult D. 2004. Microorganisms resistant to free-living amoebae. Clin Microbiol Rev 17:413–33.

Griffith DE, Aksamit T, Brown-Elliott B a, Catanzaro A, Daley C, Gordin F, Holland SM, Horsburgh R, Huitt G, Iademarco MF, Iseman M, Olivier K, Ruoss S, von Reyn CF, Wallace RJ, Winthrop K. 2007. An official ATS/IDSA statement: diagnosis, treatment, and prevention of nontuberculous mycobacterial diseases. Am J Respir Crit Care Med 175:367–416. doi:10.1164/rccm.200604-571ST

Guidotti TL, Ragain L, de Haas P, van Soolingen D. 2008. Communicating with healthcare providers. J Water Health 6:s53. doi:10.2166/wh.2008.032

Gutierrez MC, Brisse S, Brosch R, Fabre M, Omaïs B, Marmiesse M, Supply P, Vincent V. 2005. Ancient Origin and Gene Mosaicism of the Progenitor of Mycobacterium tuberculosis. PLoS Pathog 1:e5. doi:10.1371/journal.ppat.0010005

Hartman T, Weinrick B, Vilchèze C, Berney M, Tufariello J, Cook GM, Jacobs WR, Jr. 2014. Succinate dehydrogenase is the regulator of respiration in Mycobacterium tuberculosis. PLoS Pathog 10:e1004510. doi:10.1371/journal.ppat.1004510

Jamet S, Quentin Y, Coudray C, Texier P, Laval F, Daffé M, Fichant G, Cam K. 2015. Evolution of Mycolic Acid Biosynthesis Genes and Their Regulation during Starvation in Mycobacterium tuberculosis. J Bacteriol 197:3797–811. doi:10.1128/JB.00433-15

Kumar A, Farhana A, Guidry L, Saini V, Hondalus M, Steyn AJC. 2011. Redox homeostasis in mycobacteria: the key to tuberculosis control? Expert Rev Mol Med 13:e39. doi:10.1017/S1462399411002079

Laencina L, Dubois V, Le Moigne V, Viljoen A, Majlessi L, Pritchard J, Bernut A, Piel L, Roux A-L, Gaillard J-L, Lombard B, Loew D, Rubin EJ, Brosch R, Kremer L, Herrmann J-L, Girard-Misguich F. 2018. Identification of genes required for Mycobacterium abscessus growth in vivo with a prominent role of the ESX-4 locus. Proc Natl Acad Sci 115:E1002–E1011. doi:10.1073/pnas.1713195115

Lamrabet O, Medie FM, Drancourt M. 2012. Acanthamoeba polyphaga-enhanced growth of Mycobacterium smegmatis. PLoS One 7. doi:10.1371/journal.pone.0029833

Lavania M, Katoch K, Katoch VM, Gupta AK, Chauhan DS, Sharma R, Gandhi R, Chauhan V, Bansal G, Sachan P, Sachan S, Yadav VS, Jadhav R. 2008. Detection of viable Mycobacterium leprae in soil samples: Insights into possible sources of transmission of leprosy. Infect Genet Evol 8:627–631. doi:10.1016/j.meegid.2008.05.007

Le Moigne V, Belon C, Goulard C, Accard G, Bernut A, Pitard B, Gaillard J-L, Kremer L, Herrmann J-L, Blanc-Potard A-B. 2016. MgtC as a Host-Induced Factor and Vaccine Candidate against Mycobacterium abscessus Infection. Infect Immun 84:2895–2903. doi:10.1128/IAI.00359-16

Love MI, Huber W, Anders S. 2014. Moderated estimation of fold change and dispersion for RNA-seq data with DESeq2. Genome Biol 15:550. doi:10.1186/s13059-014-0550-8

Manca C, Paul S, Barry CE, Freedman VH, Kaplan G, Kaplan G. 1999. Mycobacterium tuberculosis catalase and peroxidase activities and resistance to oxidative killing in human monocytes in vitro. Infect Immun 67:74–9.

Medjahed H, Singh AK. 2010. Genetic manipulation of Mycobacterium abscessus. Curr Protoc Microbiol Chapter 10:Unit 10D.2. doi:10.1002/9780471729259.mc10d02s18

Mukhopadhyay S, Nair S, Ghosh S. 2012. Pathogenesis in tuberculosis: transcriptomic approaches to unraveling virulence mechanisms and finding new drug targets. FEMS Microbiol Rev 36:463–485. doi:10.1111/j.1574-6976.2011.00302.x

Müller MP, Peters H, Blümer J, Blankenfeldt W, Goody RS, Itzen A. 2010. The Legionella effector protein DrrA AMPylates the membrane traffic regulator Rab1b. Science 329:946–9. doi:10.1126/science.1192276

Neckers L, Tatu U. 2008. Molecular chaperones in pathogen virulence: emerging new targets for therapy. Cell Host Microbe 4:519–27. doi:10.1016/j.chom.2008.10.011

Owens CP, Chim N, Graves AB, Harmston CA, Iniguez A, Contreras H, Liptak MD, Goulding CW. 2013. The Mycobacterium tuberculosis secreted protein Rv0203 transfers heme to membrane proteins MmpL3 and MmpL11. J Biol Chem 288:21714–21728. doi:10.1074/jbc.M113.453076

Pagnier I, Raoult D, La Scola B. 2008. Isolation and identification of amoeba-resisting bacteria From water in human environment by using an Acanthamoeba polyphaga co-culture procedure. Environ Microbiol 10:1135–1144. doi:10.1111/j.1462-2920.2007.01530.x

Parra J, Marcoux J, Poncin I, Canaan S, Herrmann JL, Nigou J, Burlet-Schiltz O, Rivière M. 2017. Scrutiny of Mycobacterium tuberculosis 19 kDa antigen proteoforms provides new insights in the lipoglycoprotein biogenesis paradigm. Sci Rep 7:43682. doi:10.1038/srep43682

Rhee KY, Erdjument-Bromage H, Tempst P, Nathan CF. 2005. S-nitroso proteome of Mycobacterium tuberculosis: Enzymes of intermediary metabolism and antioxidant defense. Proc Natl Acad Sci 102:467–472. doi:10.1073/pnas.0406133102

Ribet D, Cossart P. 2010. Pathogen-Mediated Posttranslational Modifications: A Re-emerging Field. Cell 143:694–702. doi:10.1016/J.CELL.2010.11.019

Ripoll F, Pasek S, Schenowitz C, Dossat C, Barbe V, Rottman M, Macheras E, Heym B, Herrmann J-L, Daffé M, Brosch R, Risler J-L, Gaillard J-L. 2009. Non mycobacterial virulence genes in the genome of the emerging pathogen Mycobacterium abscessus. PLoS One 4:e5660. doi:10.1371/journal.pone.0005660

Rohmer M, Knani M, Simonin P, Sutter B, Sahm H. 1993. Isoprenoid biosynthesis in bacteria: a novel pathway for the early steps leading to isopentenyl diphosphate. Biochem J 295 (Pt 2:517–24.

Rossier O, Cianciotto NP. 2005. The Legionella pneumophila tatB gene facilitates secretion of phospholipase C, growth under iron-limiting conditions, and intracellular infection. Infect Immun 73:2020–32. doi:10.1128/IAI.73.4.2020-2032.2005

Roux A-L, Viljoen A, Bah A, Simeone R, Bernut A, Laencina L, Deramaudt T, Rottman M, Gaillard J-LJ-L, Majlessi L, Brosch R, Girard-Misguich F, Vergne I, de Chastellier C, Kremer L, Herrmann J-LJ-L. 2016. The distinct fate of smooth and rough Mycobacterium abscessus variants inside macrophages. Open Biol 6:160185. doi:10.1098/rsob.160185

Salah IB, Ghigo E, Drancourt M. 2009. Free-living amoebae, a training field for macrophage resistance of mycobacteria. Clin Microbiol Infect 15:894–905. doi:10.1111/j.1469-0691.2009.03011.x

Sassetti CM, Boyd DH, Rubin EJ. 2003. Genes required for mycobacterial growth defined by high density mutagenesis. Mol Microbiol 48:77–84.

Schnappinger D, Ehrt S, Voskuil MI, Liu Y, Mangan J a, Monahan IM, Dolganov G, Efron B, Butcher PD, Nathan C, Schoolnik GK. 2003a. Transcriptional Adaptation of Mycobacterium tuberculosis within macrophages: Insights into the Phagosomal Environment. J Exp Med 198:693–704. doi:10.1084/jem.20030846

Schnappinger D, Ehrt S, Voskuil MI, Liu Y, Mangan JA, Monahan IM, Dolganov G, Efron B, Butcher PD, Nathan C, Schoolnik GK. 2003b. Transcriptional Adaptation of Mycobacterium tuberculosis within Macrophages. J Exp Med 198:693–704. doi:10.1084/jem.20030846

Shah P, Swiatlo E. 2008. MicroReview A multifaceted role for polyamines in bacterial pathogens. doi:10.1111/j.1365-2958.2008.06126.x

Sherman DR, Mdluli K, Hickey MJ, Barry CE, Stover CK. 1999. AhpC, oxidative stress and drug resistance in Mycobacterium tuberculosis. Biofactors 10:211–7.

Sherman DR, Voskuil M, Schnappinger D, Liao R, Harrell MI, Schoolnik GK. 2001. Regulation of the Mycobacterium tuberculosis hypoxic response gene encoding alpha-crystallin. Proc Natl Acad Sci U S A 98:7534–9. doi:10.1073/pnas.121172498

Shin D-M, Jeon B-Y, Lee H-M, Jin HS, Yuk J-M, Song C-H, Lee S-H, Lee Z-W, Cho S-N, Kim J-M, Friedman RL, Jo E-K. 2010. Mycobacterium tuberculosis Eis Regulates Autophagy, Inflammation, and Cell Death through Redox-dependent Signaling. PLoS Pathog 6:e1001230. doi:10.1371/journal.ppat.1001230

Siddiqui R, Khan NA. 2012. Acanthamoeba is an evolutionary ancestor of macrophages: a myth or reality? Exp Parasitol 130:95–7. doi:10.1016/j.exppara.2011.11.005

Simeone R, Sayes F, Song O, Gröschel MI, Brodin P, Brosch R, Majlessi L. 2015. Cytosolic access of Mycobacterium tuberculosis: critical impact of phagosomal acidification control and demonstration of occurrence in vivo. PLoS Pathog 11:e1004650. doi:10.1371/journal.ppat.1004650

Sousa S, Bandeira M, Carvalho PA, Duarte A, Jordao L. 2015. Nontuberculous mycobacteria pathogenesis and biofilm assembly. Int J Mycobacteriology 4:36–43. doi:10.1016/J.IJMYCO.2014.11.065

Stinear TP, Seemann T, Harrison PF, Jenkin GA, Davies JK, Johnson PDR, Abdellah Z, Arrowsmith C, Chillingworth T, Churcher C, Clarke K, Cronin A, Davis P, Goodhead I, Holroyd N, Jagels K, Lord A, Moule S, Mungall K, Norbertczak H, Quail MA, Rabbinowitsch E, Walker D, White B, Whitehead S, Small PLC, Brosch R, Ramakrishnan L, Fischbach MA, Parkhill J, Cole ST. 2008. Insights from the complete genome sequence of Mycobacterium marinum on the evolution of Mycobacterium tuberculosis. Genome Res 18:729–741. doi:10.1101/gr.075069.107

Tatusov RL, Galperin MY, Natale DA, Koonin E V. 2000. The COG database: a tool for genome-scale analysis of protein functions and evolution. Nucleic Acids Res 28:33–6.

Thomas V, Herrera-rimann K, Blanc DS, Greub G. 2006. Biodiversity of Amoebae and Amoeba-Resisting Bacteria in a Hospital Water Network. Appl Env Microbiol 72:2428– 2438. doi:10.1128/AEM.72.4.2428

Thomas V, McDonnell G. 2007. Relationship between mycobacteria and amoebae: ecological and epidemiological concerns. Lett Appl Microbiol 45:349–57. doi:10.1111/j.1472-765X.2007.02206.x

Thomson RM, Carter R, Tolson C, Coulter C, Huygens F, Hargreaves M. 2013. Factors associated with the isolation of Nontuberculous mycobacteria (NTM) from a large municipal water system in Brisbane, Australia. BMC Microbiol 13:89. doi:10.1186/1471-2180-13-89

Vallenet D, Engelen S, Mornico D, Cruveiller S, Fleury L, Lajus A, Rouy Z, Roche D, Salvignol G, Scarpelli C, Médigue C. 2009. MicroScope: a platform for microbial genome annotation and Comparative genomics. Database (Oxford) 2009:bap021. doi:10.1093/database/bap021

Viljoen A, Dubois V, Girard-Misguich F, Blaise M, Herrmann J-L, Kremer L. 2017. The diverse family of MmpL transporters in mycobacteria: from regulation to antimicrobial developments. Mol Microbiol 104. doi:10.1111/mmi.13675

White CI, Birtles RJ, Wigley P, Jones PH. 2010. Mycobacterium avium subspecies paratuberculosis in free-living amoebae isolated from fields not used for grazing. Vet Rec 166:401–402. doi:10.1136/vr.b4797

Wright CC, Hsu FF, Arnett E, Dunaj JL, Davidson PM, Pacheco SA, Harriff MJ, Lewinsohn DM, Schlesinger LS, Purdy GE. 2017. The Mycobacterium tuberculosis MmpL11 Cell Wall Lipid Transporter Is important for Biofilm formation, Intracellular Growth, and Nonreplicating Persistence. Infect Immun 85:e00131–17. doi:10.1128/IAI.00131-17

Wu S, Barnes PF, Samten B, Pang X, Rodrigue S, Ghanny S, Soteropoulos P, Gaudreau L, Howard ST. 2009. Activation of the eis gene in a W-Beijing strain of Mycobacterium tuberculosis correlates with increased SigA levels and enhanced intracellular growth. Microbiology 155:1272–81. doi:10.1099/mic.0.024638-0

Yamaryo-Botte Y, Rainczuk AK, Lea-Smith DJ, Brammananth R, van der Peet PL, Meikle P, Ralton JE, Rupasinghe TWT, Williams SJ, Coppel RL, Crellin PK, McConville MJ. 2015. Acetylation of Trehalose Mycolates Is Required for Efficient MmpL-Mediated Membrane Transport in Corynebacterineae. ACS Chem Biol 10:734–746. doi:10.1021/cb5007689

