## Supplementary tables and figures for "Mycobacterium abscessus virulence traits unraveled by transcriptomic profiling in amoeba and macrophages"

**Supp. Figure legends**

**Sup. Figure 1: DESeq2 statistical analyses.**

**A.** *M. abscessus* transcriptomes in *A. castellanii* 4 and 16 hpi. **B.** *M. abscessus* transcriptome in macrophages 16 hpi**. C.** *M. chelonae* transcritome in *A. castellanii* 16 hpi. Hierarchical clustering of raw data (left panel) and transcriptome heatmaps (right panel) are depicted.

**Sup. Figure 2: Comparison of *M. abscessus* transcriptomes in *A. castellanii* and in macrophages according to differentially expressed genes fold change.**

**A.** Differentially expressed genes (DEGs) were categorized according to their fold change (FC) expressed in $Log_2$. Low DEGs depict a FC under |2|, Med DEGs depict a FC between |2| and |4| and High DEGs depict a FC higher than |4|. **B.** Ratio of UP DEGs to DOWN DEGs were represented during co-cultures with A. castellanii (Ac) and macrophages (ϕ).

**Sup. Figure 3: Selection of *M. abscessus* genes highly induced in amoebae.**

*M. abscessus* highly induced genes in *A. castellanii* (Ac) were considered (n=77). Genes poorly induced in macrophages (Mϕ) (filter 1) and during *M. chelonae* co-culture with amoeba (filter 2) were conserved (n=45). FC: fold change ($Log_2$ value).

**Sup. Figure 4: Verification of knockout strains growth in culture medium and contribution to virulence in macrophages.**

**A.** KO strains growth in culture medium. The strains were cultured in 7H9 medium supplemented with glycerol 0.1% for seven days. Growth curves were obtained by measuring the cultures optical density each day. **B.** Complementation of *M. abscessus* ΔOP,_3,-4 and 6 strains in macrophages. Macrophages were infected at 10 MOI. Experiments were repeated three times in triplicates. Statistical analyses were performed with GraphPad PRISM6.

Histograms with error bars represent means ± SD. Differences between means were analyzed by ANOVA and the Tukey post-test allowing multiple comparisons to be performed. ns = non-significant. * $p<0.05$. ** $p<0.01$. *** $p<0.001$. **** $p<0.0001$.

**Sup. Figure 5: Conservation of *M. abscessus eis* loci in *Mycobacterium tuberculosis* and vice versa.**

**A.** Conservation of *M. abscessus eis1* locus in *M. tuberculosis*. **B.** Conservation of *M. abscessus eis2* locus in *M. tuberculosis*. **C.** Conservation of *M. tuberculosis eis* locus in *M. abscessus*. Bidirectional Best Hit (BBH) search was performed between *M. abscessus* and *M. tuberculosis* genomes with the Opscan software. BBHs are depicted by arrows filled with red, brown or orange. Brown arrows correspond to MmpL-encoding genes. Orange arrows correspond to MmpS-encoding genes. Greys bands link genes or groups of genes conserved in the two species.

**Sup. Figure 6: Expression of *M. abscessus eis* genes in macrophages 4 and 16 hpi.** $Eis1_{MAB}$ (left panel) and $eis2_{MAB}$ (right panel) expression in macrophages was measured twice in triplicates by quantitative-real time PCR by normalization with *sigA* housekeeping gene.

**Sup. Figure 7: Intracellular phenotypes uncontrolled by *M. abscessus eis2* genes.**

**A.** Cell death. Macrophage death following infection with *M. abscessus* was assessed with the Dead Cell Apoptosis Kit with Annexin V FITC and PI for flow cytometry. **B.** Cell autophagy was measured Premo Autophagy Tandem Sensor RFP-GFP-LC3B Kit. At least 40 cells per condition were analyzed by confocal microscopy. The number of autophagic particules per cell (left panel) and acidification of autophagosomes (right panel) was determined with the Fiji software. **C.** Phagosomal acidification was assessed as previously described (Roux *et al.*, 2016).

Macrophages were infected at 10 (C) or 50 MOI (A and B). Histograms with error bars represent means ± SD. Differences between means were analyzed by ANOVA and the Tukey post-test allowing multiple comparisons to be performed. ns = non-significant. * $p<0.05$. ** $p<0.01$. *** $p<0.001$. **** $p<0.0001$.

### Supp. Figure 1

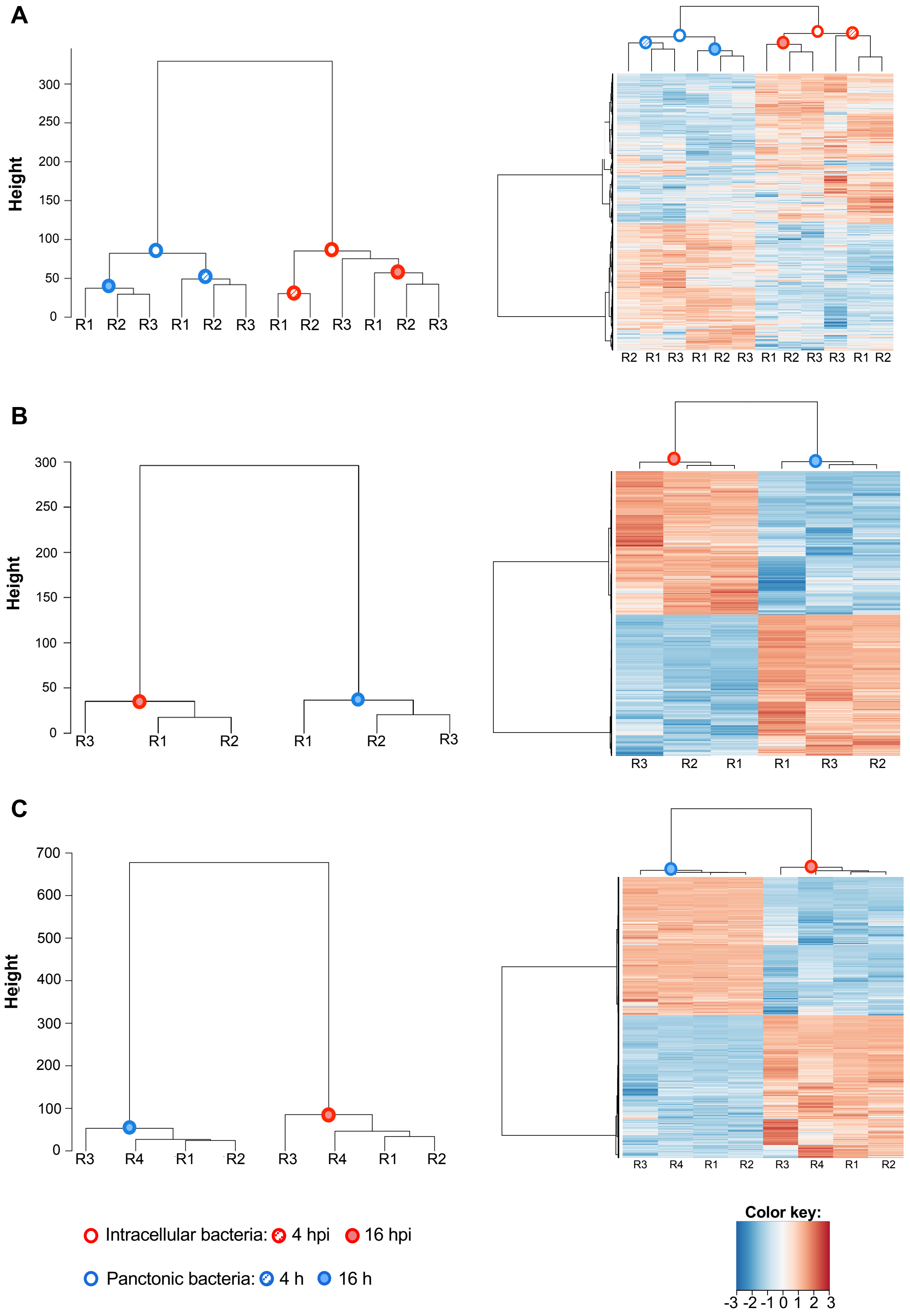

**Supp. Figure 2**

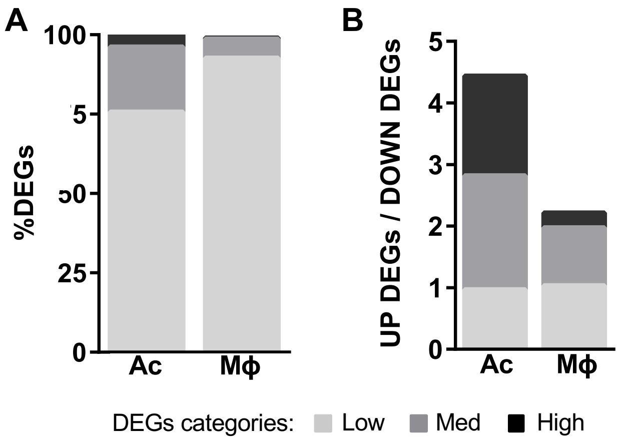

### Supp. Figure 3

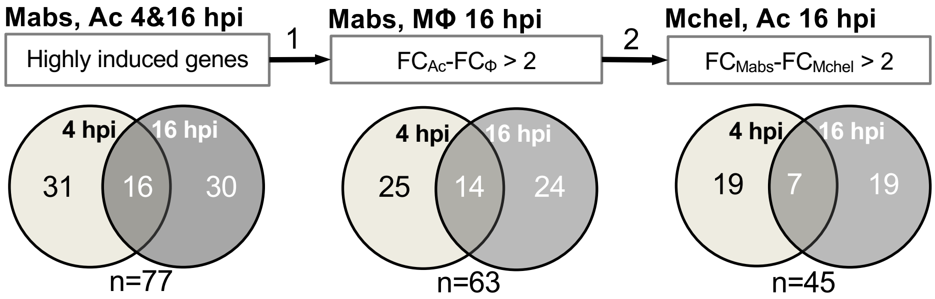

### Supp. Figure 4

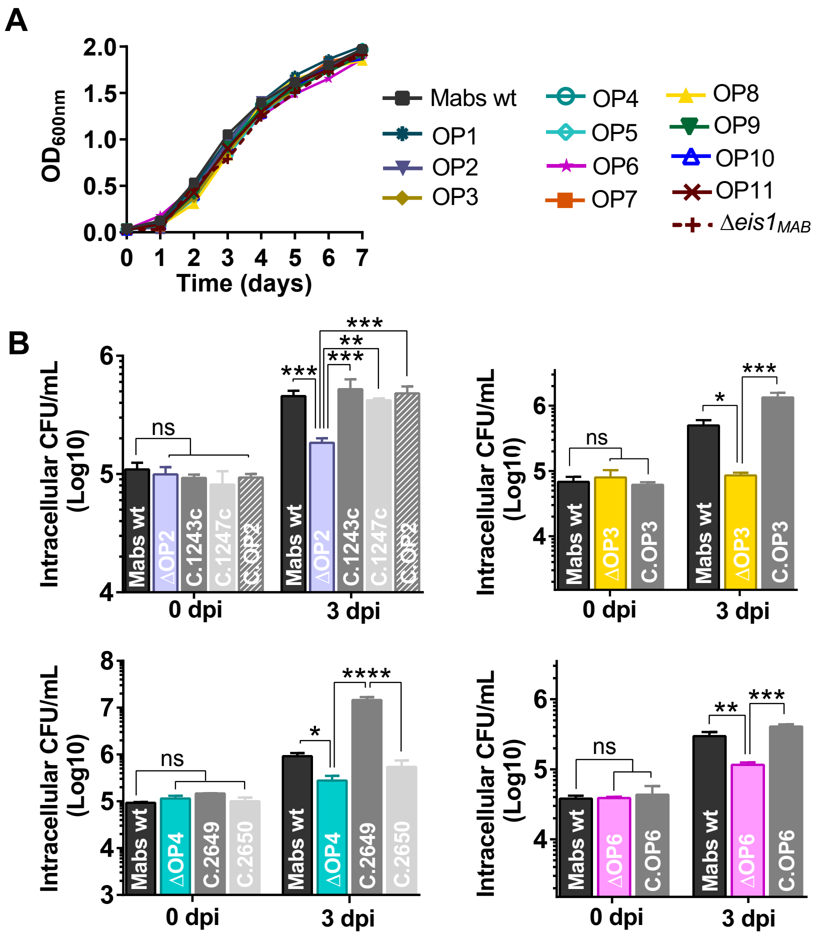

### Sup. Figure 5

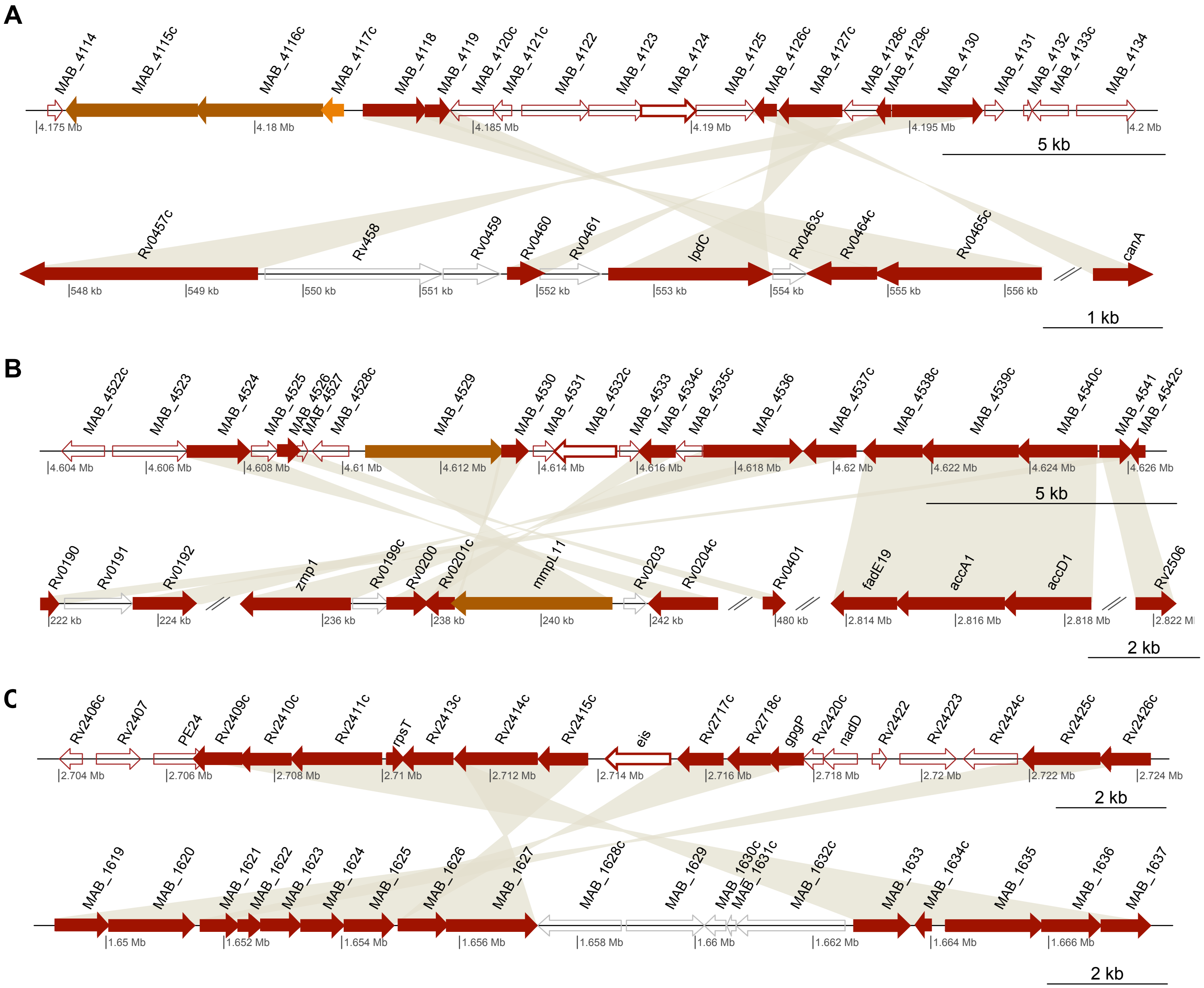
A
MAB_4114
MAB_4115c
MAB_4116c
MAB_4117c
MAB_4118
MAB_4119
MAB_4120c
MAB_4121c
MAB_4122
MAB_4123
MAB_4124
MAB_4125
MAB_4126c
MAB_4127c
MAB_4128c
MAB_4129c
MAB_4130
MAB_4131
MAB_4132
MAB_4133c
MAB_4134
4.175 Mb
4.18 Mb
4.185 Mb
4.19 Mb
4.195 Mb
4.2 Mb
5 kb
Rv0457c
Rv458
Rv0459
Rv0460
Rv0461
lpdC
Rv0463c
Rv0464c
Rv0465c
canA
548 kb
549 kb
550 kb
551 kb
552 kb
553 kb
554 kb
555 kb
556 kb
1 kb
B
MAB_4522c
MAB_4523
MAB_4524
MAB_4525
MAB_4526
MAB_4527
MAB_4528c
MAB_4529
MAB_4530
MAB_4531
MAB_4532c
MAB_4533
MAB_4534c
MAB_4535c
MAB_4536
MAB_4537c
MAB_4538c
MAB_4539c
MAB_4540c
MAB_4541
MAB_4542c
4.604 Mb
4.606 Mb
4.608 Mb
4.61 Mb
4.612 Mb
4.614 Mb
4.616 Mb
4.618 Mb
4.62 Mb
4.622 Mb
4.624 Mb
4.626 Mb
5 kb
Rv0190
Rv0191
Rv0192
zmp1
Rv0199c
Rv0200
Rv0201c
mmpL11
Rv0203
Rv0204c
Rv0401
fadE19
accA1
accD1
Rv2506
222 kb
224 kb
236 kb
238 kb
240 kb
242 kb
480 kb
2.814 Mb
2.816 Mb
2.818 Mb
2.822 Mb
2 kb
C
Rv2406c
Rv2407
PE24
Rv2409c
Rv2410c
Rv2411c
rpsT
Rv2413c
Rv2414c
Rv2415c
eis
Rv2717c
Rv2718c
gpgP
Rv2420c
nadD
Rv2422
Rv24223
Rv2424c
Rv2425c
Rv2426c
2.704 Mb
2.706 Mb
2.708 Mb
2.71 Mb
2.712 Mb
2.714 Mb
2.716 Mb
2.718 Mb
2.72 Mb
2.722 Mb
2.724 Mb
2 kb
MAB_1619
MAB_1620
MAB_1621
MAB_1622
MAB_1623
MAB_1624
MAB_1625
MAB_1626
MAB_1627
MAB_1628c
MAB_1629
MAB_1630c
MAB_1631c
MAB_1632c
MAB_1633
MAB_1634c
MAB_1635
MAB_1636
MAB_1637
1.65 Mb
1.652 Mb
1.654 Mb
1.656 Mb
1.658 Mb
1.66 Mb
1.662 Mb
1.664 Mb
1.666 Mb
2 kb

### Supp. Figure 6

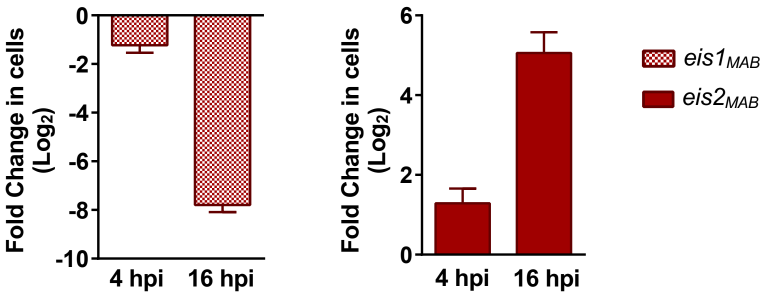

### Supp. Figure 7

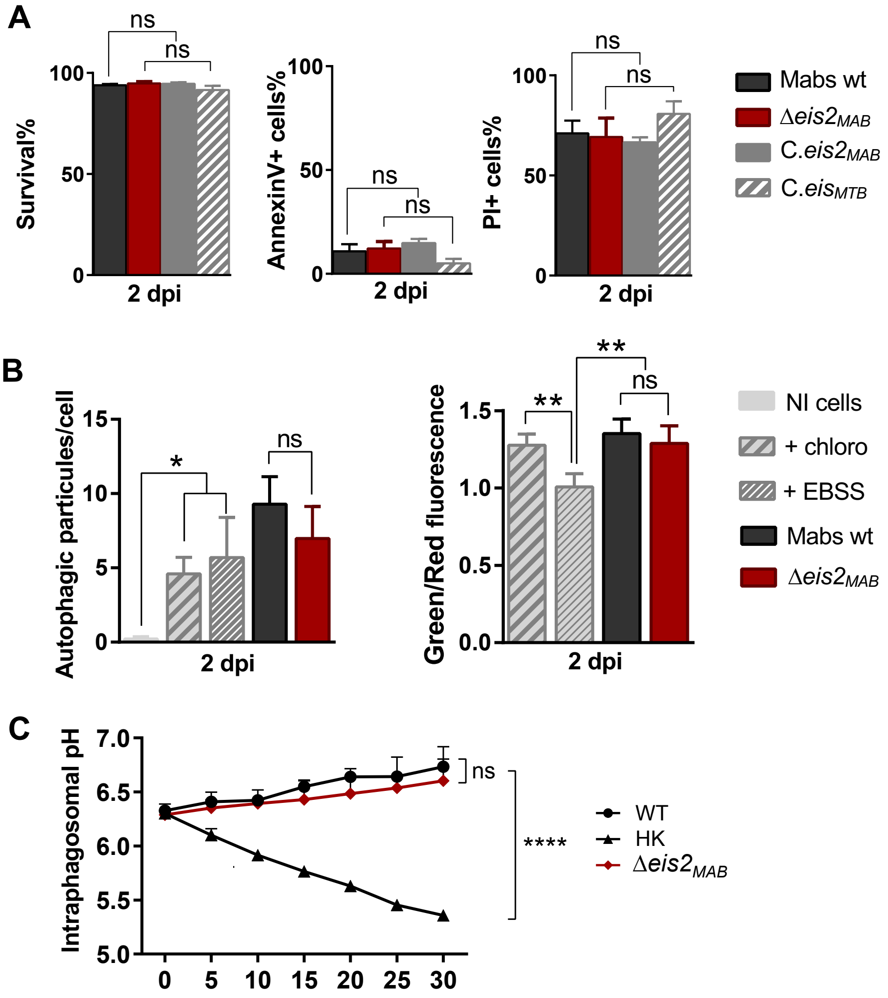

**Supp. Table 1: Differentially expressed genes identified with the *DEseq2* package.**

| Transcriptomes | Down DEGs | Up DEGs | Tot | CDS cover |
|---|---|---|---|---|
| Mabs, Ac 4 hpi | 1218 | 1355 | 2573 | 56% |
| Mabs, Ac 16 hpi | 1095 | 1122 | 2217 | 49% |
| Mabs, Mϕ hpi | 1646 | 1671 | 3317 | 73% |
| Mchel, Ac 16 hpi | 1186 | 1949 | 3835 | 80% |

**Supp. Table 2: List of *M. abscessus* genes highly induced in Ac only.**

| Mma gene | Encoded protein | IP5 analysis (IPR) | Mabs BBH | MTB BBH | MAV BBH | FC Ac, 4 hpi | FC Ac, 16 hpi |
|---|---|---|---|---|---|---|---|
| ***Highly induced genes 4 hpi*** | | | | | | | |
| *MYCMA_RS01100* | Hypothetical protein | No IPR | *MAB_4664* | | | 5.71 | 0.00 |
| *MYCMA_RS08980* | Glyoxalase | | *MAB_3056c* | | | 5.32 | 1.81 |
| *MYCMA_RS10865* | Hypothetical protein | Magnesium transporter MgtE (006669) | *MAB_2717c* | | *MAV_2122* | 5.26 | 1.44 |
| *MYCMA_RS14505* | MULTISPECIES: 3-methyl-2-oxobutanoate hydroxymethyltransferase | | *MAB_1916c* | *panB* | *panB* | 5.22 | 1.89 |
| *MYCMA_RS17210* | Copper oxidase | | *MAB_1271c* | | | 4.99 | 2.71 |
| *MYCMA_RS10860* | Hypothetical protein | No IPR | - | | | 4.99 | 1.26 |
| *MYCMA_RS11360* | Monooxygenase | | *MAB_2607* | | | 4.93 | 0.00 |
| *MYCMA_RS10755* | Hypothetical protein | No IPR | *MAB_2738c* | | | 4.84 | 2.22 |
| *MYCMA_RS17970* | Hypothetical protein | No IPR | *MAB_1117c* | | | 4.49 | 2.33 |
| *MYCMA_RS07655* | MULTISPECIES: alkylhydroperoxidase | | *MAB_3311c* | | | 4.48 | 3.25 |
| *MYCMA_RS02005* | Short-chain dehydrogenase | | *MAB_4478c* | | *MAV_1797* | 4.45 | 1.84 |
| *MYCMA_RS06830* | Nitroreductase | | *MAB_3461c* | | | 4.38 | 2.11 |
| *MYCMA_RS20755* | Membrane protein | Transport accessory protein MmpS (008693) | *MAB_0477* | | | 4.31 | 2.77 |
| *MYCMA_RS09135* | Hypothetical protein | Virulence factor BrkB (017039) | *MAB_3025* | *Rv2707* | *MAV_3600* | 4.29 | 1.90 |
| *MYCMA_RS11655* | Hypothetical protein | No IPR | - | | | 4.23 | 1.87 |
| *MYCMA_RS08985* | Pyridoxamine 5'-phosphate oxidase | | *MAB_3055c* | | | 4.16 | 0.00 |
| *MYCMA_RS11985* | Sulfite reductase subunit alpha | | *MAB_2492* | | | 4.13 | 2.55 |
| *MYCMA_RS12840* | Hypothetical protein | No IPR | *MAB_2313* | | | 4.07 | 3.46 |
| *MYCMA_RS03450* | Acyl-CoA dehydrogenase | | *MAB_4158* | *fadE26* | *MAV_0652* | 4.05 | 2.31 |
| ***Highly induced genes 4 & 16 hpi*** | | | | | | | |
| *MYCMA_RS18820* | Sugar translocase | | - | | | 6.52 | 6.22 |
| *MYCMA_RS11145* | MULTISPECIES: MmpS family protein | | *MAB_2649* | | | 4.73 | 6.82 |

| | | | | | | | |
|---|---|---|---|---|---|---|---|
| *MYCMA_RS19885* | Short-chain dehydrogenase | | *MAB_0646c* | *Rv0068* | *MAV_4710* | 5.67 | 5.27 |
| *MYCMA_RS01105* | Hypothetical protein | No IPR | *MAB_4663* | | | 6.16 | 4.18 |
| *MYCMA_RS00455* | Hypothetical protein | Twin-arginine translocation pathway, signal sequence (006311) | *MAB_4791c* | | | 5.17 | 4.96 |
| *MYCMA_RS21900* | TQXA domain-containing protein | | *MAB_0219* | | *MAV_2053* | 5.33 | 4.73 |
| *MYCMA_RS07540* | MULTISPECIES: ATP-binding protein | | *MAB_3325c* | | | 5.03 | 4.75 |
| ***Highly induced genes 16 hpi*** | | | | | | | |
| *MYCMA_RS13965* | (Fe-S)-cluster assembly protein | | *MAB_2020c* | | | 2.77 | 4.93 |
| *MYCMA_RS17325* | Hypothetical protein | No IPR | *MAB_1244c* | | | 3.11 | 5.03 |
| *MYCMA_RS06800* | MerR family transcriptional regulator | | - | | | 2.71 | 4.00 |
| *MYCMA_RS15995* | TetR family transcriptional regulator | | *MAB_1518* | | *MAV_4046* | 1.52 | 4.01 |
| *MYCMA_RS18825* | MarR family transcriptional regulator | | *MAB_0925c* | | | 3.20 | 5.04 |
| *MYCMA_RS17330* | Hypothetical protein | ABC transporter, FecCD/TroCD-like | *MAB_1243c* | | | 2.86 | 4.97 |
| *MYCMA_RS17315* | Hypothetical protein | Alkaline shock protein Asp23 (05531) | *MAB_1247c* | | | 3.43 | 5.10 |
| *MYCMA_RS02820* | MULTISPECIES: molecular chaperone | | *MAB_4273c* | *dnaK* | *dnaK* | 2.63 | 4.87 |
| *MYCMA_RS12505* | Activator of HSP90 ATPase | | *MAB_2387* | | | 3.12 | 4.99 |
| *MYCMA_RS20640* | Carboxymuconolactone decarboxylase | | - | | | 0.00 | 7.63 |
| *MYCMA_RS10645* | YrbE family protein | | - | | | 0.00 | 4.71 |
| *MYCMA_RS16000* | GlcNAc transferase | | *MAB_1517c* | | *MAV_4048* | 0.00 | 4.31 |
| *MYCMA_RS17320* | Hypothetical protein | No IPR | *MAB_1246c* | | | 2.07 | 4.27 |
| *MYCMA_RS09170* | GntR family transcriptional regulator | | *MAB_3018* | *Rv0586* | *MAV_4554* | 0.00 | 5.71 |
| *MYCMA_RS10650* | Membrane protein | No IPR | - | | | 0.00 | 4.17 |
| *MYCMA_RS01300* | TetR family transcriptional regulator | | *MAB_4625* | | | 2.55 | 4.30 |
| *MYCMA_RS10655* | Membrane protein | No IPR | - | | | 0.00 | 4.05 |
| *MYCMA_RS02345* | MULTISPECIES: alkyl hydroperoxide | | *MAB_4408c* | *ahpC* | *MAV_2839* | 3.46 | 4.04 |
| *MYCMA_RS14940* | Transposase | | - | | | 3.22 | 4.03 |

**Supp. Table 3 : List of *M. abscessus* genes highly induced in M**ϕ or Ac 16 hpi.

| Mma gene | Encoded protein | IP5 analysis (IPR) | Mabs gene | MTB BBH | MAV BBH | FC Mϕ | FC Ac |
|---|---|---|---|---|---|---|---|
| *MYCMA_RS01880* | Hypothetical protein | No IPR | *MAB_4509c* | - | - | 5.78 | 5.11 |
| *MYCMA_RS01765* | Hypothetical protein | N-acetyltransferase Eis (016181) | *MAB_4532c* | - | - | 5.37 | 2.09 |
| *MYCMA_RS13035* | MFS transporter | | *MAB_2273* | - | - | 5.26 | 4.15 |
| *MYCMA_RS17085* | Acyltransferase | | *MAB_1297c* | - | *MAV_4113* | 4.86 | 4.91 |
| *MYCMA_RS08590* | Transcriptional regulator | | *MAB_3134c* | - | - | 4.54 | 1.55 |
| *MYCMA_RS08595* | Hemin transporter | | *MAB_3133c* | - | - | 4.36 | 1.14 |
| *MYCMA_RS06600* | MULTISPECIES: transcriptional regulator | | *MAB_3508c* | *whiB7* | *MAV_4142* | 4,24 | 4,37 |
| *MYCMA_RS02565* | Acetyltransferase | | *MAB_4324c* | - | - | 4.18 | 0.00 |
| *MYCMA_RS05440* | Membrane protein | EamA domain (000620) | *MAB_3762* | - | - | 3.96 | 4.49 |
| *MYCMA_RS12630* | ABC transporter | | *MAB_2355c* | - | - | 3.92 | 1.76 |
| *MYCMA_RS17930* | GNAT family acetyltransferase | | *MAB_1125c* | - | - | 3.89 | 1.62 |
| *MYCMA_RS05315* | Hypothetical protein | No IPR | *MAB_3786c* | - | - | 3.85 | 0.90 |
| *MYCMA_RS08600* | Membrane protein | No IPR | *MAB_3132c* | *Rv2620c* | *MAV_3498* | 3.83 | 0.00 |
| *MYCMA_RS19730* | Membrane protein | EamA domain (000620) | *MAB_0677c* | - | - | 3.83 | 0.00 |
| *MYCMA_RS22615* | Esterase | | *MAB_0078* | - | *MAV_3025* | 3.82 | 2.92 |
| *MYCMA_RS13590* | DEAD/DEAH box helicase | | *MAB_2158c* | - | *MAV_2956* | 3.81 | 4.38 |
| *MYCMA_RS11580* | Universal stress protein UspA | | - | - | - | 3.78 | 0.00 |
| *MYCMA_RS06595* | Hypothetical protein | No IPR | *MAB_3509c* | - | - | 3.72 | 3.35 |
| *MYCMA_RS22620* | IclR family transcriptional regulator | | *MAB_0077* | - | *MAV_3024* | 3.69 | 2.54 |
| *MYCMA_RS08355* | Membrane protein | EamA domain (000620) | *MAB_3180* | - | *MAV_0095* | 3.68 | -1.60 |
| *MYCMA_RS04790* | DNA-binding response regulator | | *MAB_3891c* | *devR* | *MAV_4109* | 3.62 | -1.02 |
| *MYCMA_RS19020* | Hypothetical protein | FAD//NAD(P)-binding domain superfamily (I036188) | *MAB_0857* | - | - | 3.55 | 2.73 |
| *MYCMA_RS10510* | Transporter | | *MAB_2780c* | - | - | 3.47 | 0.83 |
| *MYCMA_RS13570* | Hypothetical protein | No IPR | - | - | | 3.45 | 0.00 |
| *MYCMA_RS16570* | MFS transporter | | *MAB_1409c* | *Rv1258c* | *MAV_1406* | 3.44 | 2.59 |
| *MYCMA_RS22575* | Taurine catabolism dioxygenase | | *MAB_0086* | *Rv3406* | - | 3.41 | 5.90 |
| *MYCMA_RS19295* | Membrane protein | | *MAB_0766* | - | - | 3.36 | 0.00 |
| *MYCMA_RS11690* | Transcriptional regulator | | *MAB_2562c* | *Rv0081* | *MAV_5108* | 3.32 | 0.00 |
| *MYCMA_RS21395* | Catalase | | *MAB_0351* | - | *katA* | 3.27 | 0.00 |
| *MYCMA_RS00530* | SAM-dependent methyltransferase | | - | - | - | 3.26 | 0.00 |

| | | | | | | | |
|---|---|---|---|---|---|---|---|
| *MYCMA_RS07025* | Hypothetical protein | 2 isopropylmate synthase LeuA, allosteric (dimerization) domain superfamily (036230) | *MAB_3424c* | - | *MAV_3928* | 3.25 | 3.06 |
| *MYCMA_RS19455* | Hypothetical protein | No IPR | *MAB_0733* | - | - | 3.22 | 1.62 |
| *MYCMA_RS06265* | Alkane 1-monooxygenase | | *MAB_3598c* | *alkB* | *MAV_4215* | 3.14 | 2.51 |
| *MYCMA_RS15610* | Guanylate cyclase | Papain-like cysteine peptidase superfamily (IPR038765) | *MAB_1591* | *Rv1118c* | *MAV_1249* | 3.14 | 0.00 |
| *MYCMA_RS14880* | ABC transporter | | *MAB_1846* | - | - | 3.09 | 2.68 |
| *MYCMA_RS14875* | Peptidase | | *MAB_1847* | - | - | 3.04 | 0.87 |
| *MYCMA_RS02705* | MULTISPECIES: aminotransferase AlaT | | *MAB_4294* | *aspC* | *MAV_4818* | 3.04 | 1.54 |
| *MYCMA_RS19450* | Hypothetical protein | Leukocidin/porin MspA superfamily (036435) | *MAB_0734* | - | - | 3.00 | 3.15 |
